# Planar polarization of cilia in the zebrafish floor-plate involves Par3-mediated posterior localization of highly motile basal bodies

**DOI:** 10.1101/702282

**Authors:** Antoine Donati, Sylvie Schneider-Maunoury, Christine Vesque

## Abstract

To produce a directional flow, ciliated epithelia display a uniform orientation of ciliary beating. Oriented beating requires planar cell polarity (PCP), which leads to planar orientation and asymmetric positioning of the ciliary basal body (BB) along the polarity axis. We took advantage of the polarized mono-ciliated epithelium of the embryonic zebrafish floor plate to investigate by live-imaging the dynamics and mechanisms of BB polarization. We showed that BBs, although bearing a cilium, were highly motile along the antero-posterior axis. BBs contacted both the anterior and the posterior membranes, with a bias towards posterior contacts from early somitogenesis on. Contacts exclusively occurred at junctional Par3 local enrichments or “patches” and were often preceded by transient membrane digitations extending towards the BB, suggesting focused cortical pulling forces. Accordingly, BBs and Par3 patches were linked by dynamic microtubules. We showed that Par3 became posteriorly enriched prior to BB posterior positioning and that floor plate polarization was impaired upon Par3 patches disruption triggered by Par3 or aPKC overexpression. In the PCP mutant *Vangl2*, where floor plate cells fail to polarize, we observed that BB were still motile but presented behavioral defects, such as ectopic contacts with lateral membranes that correlated with Par3 patch fragmentation and spreading to lateral membranes. Our data lead us to propose an unexpected function for posterior local Par3 enrichment in controlling BB asymmetric positioning downstream of the PCP pathway via a microtubule capture/shrinkage mechanism.

## INTRODUCTION

Cilia are conserved microtubule-based organelles with sensory and motile functions that are nucleated from a modified centriole called the basal body (BB). Motile cilia generate forces sufficient to propel whole organisms or bodily fluids within cavities in animals (^1, 2^). In order to generate a directional flow, ciliated epithelia display a uniform orientation of ciliary beating, which is a form of planar cell polarity (PCP). Oriented beating of a cilium usually involves two PCP processes: the off-centering of the cilium BB (translational polarity, in monociliated epithelia and ependymal cells) and the orientation of its beating relative to the main tissue axis (rotational polarity) (^1^).

In many vertebrate ciliated tissues such as the mouse cochlea and ependyma, the laterality organ of mouse and zebrafish, the Xenopus larval skin and the zebrafish floor plate, cilium polarity requires the PCP pathway. In these tissues, PCP proteins such as Van Gogh like 2 (Vangl2), Frizzled (Fz3/6), Cadherin EGF LAG seven-pass G-type receptors (Celsr1-3) and Dishevelled (Dvl1-3), localize asymmetrically in ciliated epithelia, and are required for proper cilia/BB positioning (^3,4,5,6,7,8^).

Outside the PCP pathway, the cellular and molecular mechanisms of BB positioning remain poorly understood. Non-muscle myosin II is required for ependymal translational polarity in murine ependymal multiciliated cells (^9^) and the murine Myosin Id mutant exhibit defects in both translational and rotational polarity in these cells (^10^). Translational polarity has been shown to require Rac1 in monociliated cells of the mouse node and cochlea (^11, 12^) and G protein signalling in cochlear hair cells (^13, 14^). Ciliary proteins themselves have been involved in planar polarization of cilia in several contexts (^6, 15,16,17,18^). However, the relationships between these different actors and how they impact basal body movement is unclear.

Understanding the mechanisms of cilium polarization would highly benefit from a dynamic analysis of BB movements. A major drawback is the difficulty to follow the dynamics of BB polarization *in vivo* in whole embryos, or to reproduce PCP and cilium polarization *in vitro* in cultured cells. So far, live imaging of cilium polarization has been performed only in cochlear explants where only confined Brownian motion of centrioles was observed (^19^) and in the mouse node (^11^) and ependyma (^9^) with limited temporal resolution. In this paper, in order to get a better understanding of the mechanisms leading to BB off-centering in epithelia, we have used the zebrafish embryonic floor-plate (FP) as a convenient system to investigate the dynamics of the polarization process in live embryos. The FP is a simple mono-ciliated epithelium whose posterior-positioned motile cilia allow circulation of the embryonic cerebrospinal fluid (^20^).

Our results show that planar polarization of BBs and their associated cilia is progressive during somitogenesis and is accompanied by a change in the behavior of the BBs, which are highly motile at early stages and tend to spend an increasing amount of time in contact with the posterior membrane as development proceeds. We found that BBs always contacted membranes at Par3-enriched apical junctions. Par3 became enriched at the posterior apical side of FP cells before BB polarization. Par3 and aPKC overexpression disrupted FP polarization and Par3 distribution along apical junctions was also disrupted in a *Vangl2* mutant. Thus, we propose that a major role of the PCP pathway in the FP is to drive Par3 asymmetric localization, which in turn mediates BB posterior positioning.

## RESULTS

### Floor-plate polarization shows temporal progression but no spatial synchronization

Posterior positioning of the BB in the zebrafish FP is visible as soon as 18 hours post-fertilization (hpf) (^17^) and is maintained at least until 72 hpf (^21^). From 24 hpf onward, coupled to posterior tilting of cilia, it is instrumental in propelling the cerebrospinal fluid in the spinal cord central canal (^5, 22^). At late gastrulation stages (10 hpf), ectodermal cells already display a slight posterior bias of centrioles (^23^).

To define the time-course of FP cell polarization, we assessed basal-body (BB) position along the antero-posterior (A/P) axis on fixed embryos from the 6 s to the 26 s stage (“s” stands for “somites”)(Fig. 1b, c). For each cell we defined a BB polarization index (p.i. in Fig. 1a). BBs already exhibited a posterior bias at 6 s, and the polarization state did not change significantly until 10 s. From 10 s onward, there was a progressive increase in FP polarization, mostly due to an increase in the percentage of cells with a BB in contact with the posterior membrane, with a concomitant disappearance of anterior BBs and a reduction of median BBs. The polarization state of the FP was considered complete at 18 s, since no significant difference could be detected between the 18 s and 26 s stages (Fig. 1c). Interestingly, we did not detect a gradient of polarization index along the A/P axis of the spinal cord (Fig. S1a), and single non-polarized cells were often intermingled among polarized neighbors (Fig. S1b), arguing against the existence of polarization waves.

**Figure 1.**
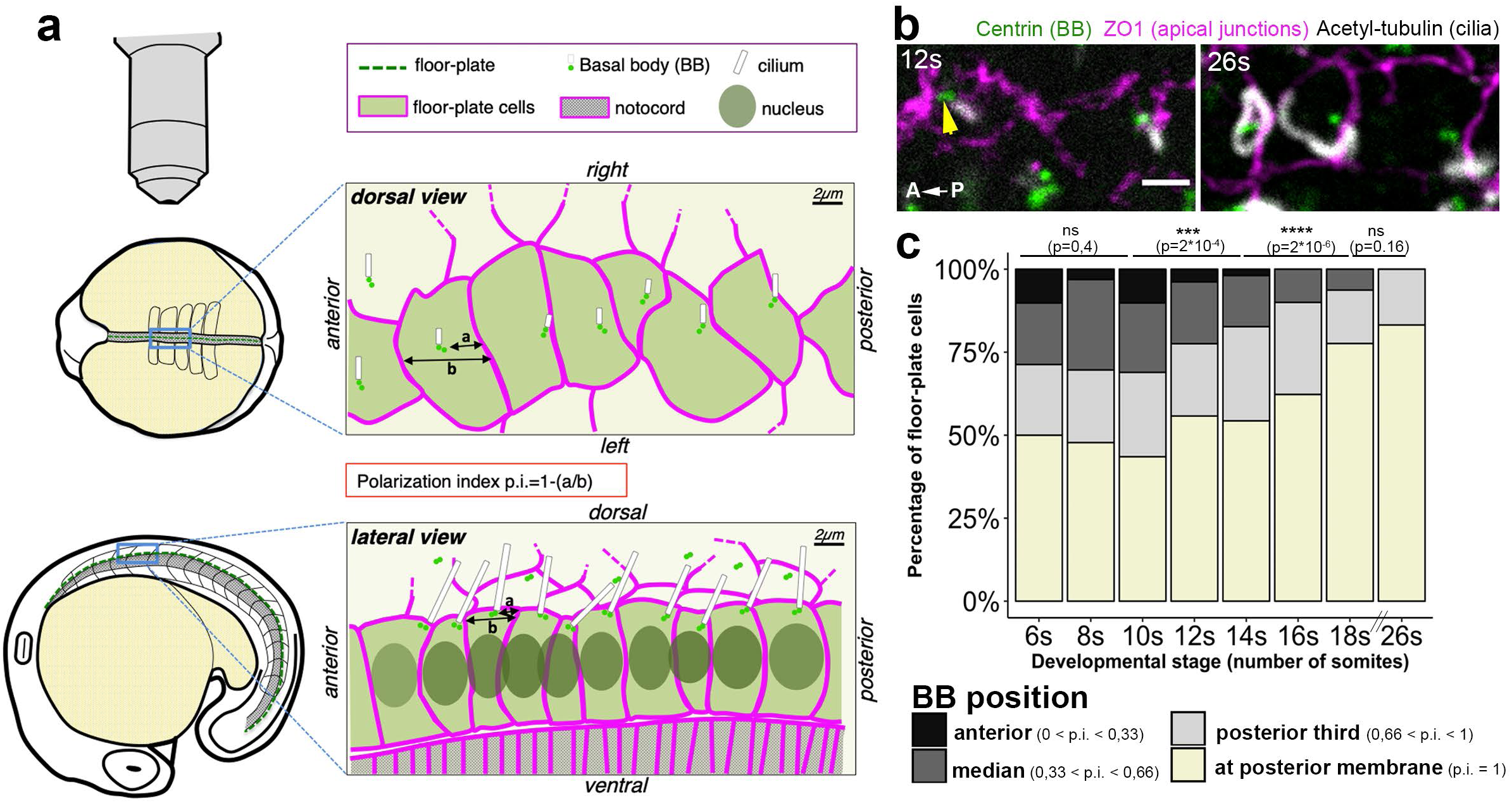
Zebrafish floor-plate progressive planar polarization during somitogenesis. **a)** Experimental set-up used to study floor-plate planar polarization in fixed or live embryos. Early stage embryo (4-12s stages) or late (after 18s) stage embryos, which displayed floor-plate cells with large apical surfaces were usually imaged from the top (dorsal view, upper part of the figure, see also b)), whereas embryos at intermediate stages (with narrower apical surfaces) were imaged from the side (lateral view, bottom half of the figure). A polarization index (defined as p.i.=1-(a/b) where “a” is the distance between the BB and the posterior membrane and “b” the distance between anterior and posterior membranes) was used to quantify BB position along the antero-posterior axis. **b-c)** Time-course of floor-plate polarization between the 6 s and 26 s stages **b)** Dorsal views of the floor-plate of flat-mounted embryos showing immunostaining against Centrin (green, BB), ZO1 (magenta, apical junctions) and Acetylated-Tubulin (white, cilia) at 12 s (left) and 26 s (right). Note that cilia are already visible at 12 s but are much longer at 26 s. The yellow arrow points at an anterior BB associated to a cilium. *Scale bar:* 2 µm **c)** Quantification of BB position measured from immuno-stained samples as shown in a. BB position along the anterior-posterior axis was quantified using the polarization index. Cells were then allocated to different categories depending on their polarization index for each developmental stage (6 s: 7embryos, 108 cells; 8 s: 14 embryos, 224 cells; 10 s: 14 embryos, 354 cells; 12 s: 5 embryos, 156 cells; 14 s: 9 embryos, 208 cells; 16 s,: 9 embryos, 220 cells; 18 s: 5 embryos, 143 cells; 26 s: 4 embryos, 119 cells). Statistical significance was assessed using a Wilcoxon test.

### BBs are highly motile in FP cells

We then turned to live-imaging to obtain a dynamic view of the polarization process. We followed BB movements within the apical surface of individual FP cells at different developmental stages (4 s to 21 s). We found that BBs displayed a highly motile behavior within the apical surface (Fig. 2a-d and Movies S1-S4), moving both anteriorly and posteriorly (Fig. S1d, first column and Fig. 6a ‘wt’). At early stages, there was a clear BB movement orientation bias along the antero-posterior axis: 70% of BB movements were oriented along this axis, whereas only 30% were oriented along the left-right axis (Fig. 6a, b, wt). However there was no overall significant difference between the number of anterior and posterior movements (although there seems to be a tendency towards more posterior movements, as seen in Fig. 6a).

**Figure 2.**
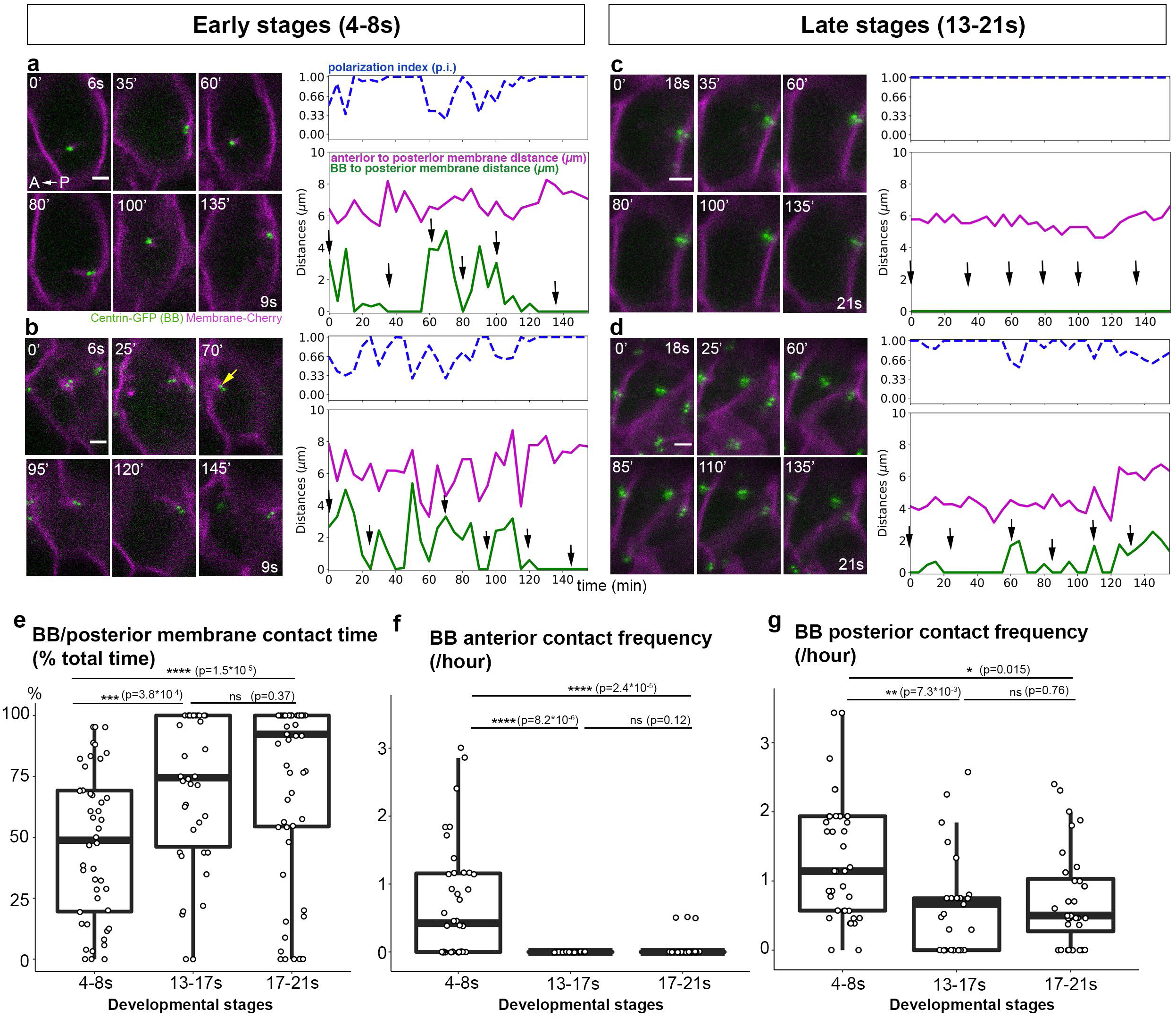
Floor-plate planar polarization involves a change in basal body (BB) motile behavior. a-d) Live imaging of BB movements during the polarization process. Images were taken every 5 minutes; a selection of images is presented here from two early stage embryos (**a, b,** movies between the 6 s and 9 s stages; **d** yellow arrow in b points at an anterior contact event) and two late stage embryos (**c, d,** movies between the 18 s and 21 s stages). The distances between BBs and posterior membranes were then plotted (green curve) along with the distance between the anterior and posterior membranes (magenta curve) and the p.i. (dashed blue curve). Black arrows on the graphs indicate the position of the images displayed on the left. **e)** Quantification of the percentage of total movie time spent by the BB in contact with the posterior membrane. (4-8s: 5 embryos, 41 cells; 13-17s: 6 embryos, 38 cells; 17-21s: 7 embryos, 59 cells). **f, g)** Number of contact events per h between BB and anterior (**f**) or posterior (**g**) membrane in embryos filmed at different developmental stages: 4 to 8 s (5 embryos, 41 cells), 13 to 17 s (5 embryos, 25 cells) and 17 to 21 s (7 embryos, 32 cells). Cells with a BB in contact with the posterior membrane during the whole movie (points at 100% in Fig. 1e) were not plotted in f and g. Statistical significance was assessed using a Wilcoxon test. *Scale bars:* 2 µm.

Cell deformations along the AP axis were more important at early stages (4-10 s) (Fig. 2a, b) than at later stages (14-21 s) (Fig. 2c, d), probably as a consequence of convergence-extension movements, but even at early stages we could see many long BB movements that did not correlate with cell deformation (see for example the two large anterior and then posterior movements around 55 and 75 min in Fig. 2a). This suggest that BBs are actively moving within FP cell apical surfaces and not just passively moved by cell deformation. One possible explanation for the presence of unpolarized cells is that they could have just undergone mitosis (during which one of the centrosomes migrates to the anterior side). However, mitoses were rare in FP cells at early stages (6 / 79 cells, 9 embryos at 4-8 s) and absent at later stages (118 cells from 15 embryos at 13-21s). Thus the impact of cell shape changes and mitosis on FP polarization is likely very small.

### FP polarization involves a change in BB behavior

In order to characterize BB behavioral changes during development, we determined the percentage of time that BBs spent in contact with the posterior membrane (Fig. 2e). At early stages, BBs spent in average 44% of their time in contact with the posterior membrane, versus more than 70% at later stages (13-21 s). This was largely due to an increase in the number of cells in which the BB stayed in contact with the posterior membrane during the whole movie (Fig. 2c). We refer to this situation as “posteriorly docked BB”. At early stages (4-8s), we did not observe any cell with posteriorly docked BB (41 cells, 5 embryos), whereas they made up 34% of the FP cell population at 13-17s stages (13/38 cells, 6 embryos) and almost half (46%) the FP population at later stages (17-21s, 27/59 cells, 7 embryos). We also noted a decrease in the frequency of BB direction changes, as well as an increase in the mean duration of BB/posterior membrane contact events and mean polarization index, suggesting that, as development proceeds, BB movements are less dynamic and more confined to the posterior side of the cell (Fig. S1d, first line). Posteriorly docked BBs made a significant contribution to these behavioral changes. In order to determine if changes in the behavior of non-posteriorly docked BBs contributed to the increase of FP polarization during somitogenesis, we quantified the same parameters, but taking into account only these motile BBs (Fig. S1d, second line): although less drastic, the same trend in BB behavior change was observed.

To further characterize the behavior of non-posteriorly docked BB, we quantified the frequency of contact events between the BB and either the anterior or the posterior membrane (Fig. 2f and g, respectively). First, posterior contacts were more frequent than anterior ones even at 4-8s (compare Fig. 2f and g), confirming that FP cells already had a posterior polarization bias at these early stages. Second, contacts with the anterior membrane were frequently observed at early stages (50% of BBs made at least one anterior contact per hour, see for example at t=70’ in Fig. 2b), but almost never observed at later stages (only 3/57 cells displayed one anterior contact). Contact frequency with the posterior membrane was also significantly higher at earlier stages (1.3 contact/h on average) than at later stages (around 0.8 contact/hour in average within the 13-21s stage window, Fig. 2g). This reduction in the number of contacts could be due to an increase in their duration (Fig. S1d, plot 2^nd^ column, 2^nd^ line) and to a reduction in BB speed. Indeed, we found that BBs moved faster at earlier stages (Fig. S1c, median speed was 0.2 µm/min at 4-8 s versus 0.1 µm/min at 13-21 s). Thus, the observed changes in FP polarization are explained both by an increase in the posteriorly docked BB population and by behavioral changes (reduced speed, less direction changes, longer posterior contact events) in other BBs.

### Transverse membrane digitations elongate towards motile floor-plate BBs

Live-imaging revealed the presence of membrane digitations extending between the BB and transverse membranes (Fig. 3; Movies S5 and S6). At early stages, we could detect such digitations in 44% of FP cells (taking into account only non-posteriorly docked BBs) (26/59 cells, 9 embryos), most of which were linking the posterior membrane and the BB (83%, 45/54 digitations, Fig. 3a upper row, white arrow, MovieS5), although digitations from the anterior membrane were also seen (Fig. 3a second row, Movie S6) (Fig. 3b). These early stage digitations were most of the time observed on a single time frame (anterior digitations) or two consecutive timeframes in time-lapse movies with a 2min or 5min time interval (Δt2min or Δt5min, respectively) between two images, but we could not detect a significant difference of digitation lifetime between anterior and posterior digitations (Fig. 3c). Posterior digitations were followed by a posterior directed BB movement in 67% of cases (26/39 digitations), whereas anterior digitations were followed by a BB anterior movement in only 22% of cases (2/9 anterior digitations) (Fig. 3d). Membrane digitations were rarely seen at later stages (after 14s, 9/40 cells, 10 embryos), probably in part because BBs spent a higher fraction of their time associated with the posterior membrane (see Fig. 1 and Fig. S1).

**Figure 3.**
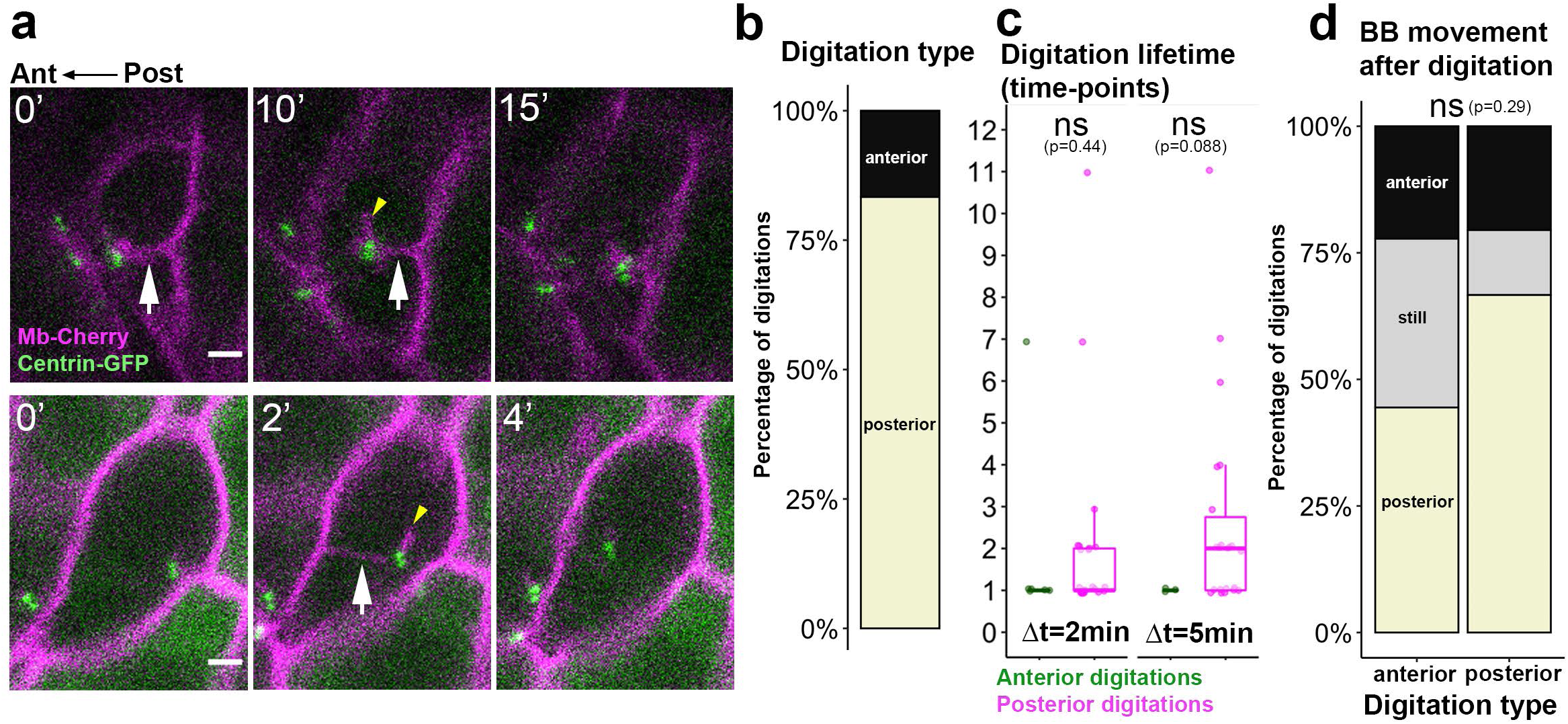
Membrane digitations link BBs to transverse membranes during FP polarization. **a)** Images taken from live-imaging (Δt = 2 or 5 min between two images) showing a posterior digitation (top) and an anterior digitation (bottom) (white arrows). Time (in min) is indicated in the upper-left corner. Short mbCherry-positive digitations, presumably corresponding to cilia, were in some cases associated to the BB (yellow arrowheads). These membrane digitations were rare in late stage embryos (6/57 cells out of 10 embryos) compared to early embryos (44/68 cells from 9 embryos), suggesting that Mb-Cherry entry into cilia is less common at later stages, which could reflect a maturation of the “ciliary gate”, a set of proteins regulating entry in and exit out of cilia. **b)** Proportion of posterior and anterior digitations (54 digitations, 8 embryos, 22 cells) **c)** Number of timepoints where anterior or posterior digitations were detected in time-lapse movies with Δt = 2 or 5 minutes (Δt2min: 6 anterior, 23 posterior digitations, 4 embryos, 8 cells. Δt5min: 3 anterior and 22 posterior digitations, 3 embryos, 13 cells, Wilcoxon tests) **d)** BB movements after anterior (left bar) or posterior digitation (48 digitations, 8 embryos, 22 cells, Fisher test)

In order to better characterize these membrane digitations at early developmental stages, we imaged the floor-plate of wt embryos at a high temporal resolution and acquired a z-stack every 10 seconds (hereafter, Δt10sec movies). Digitations were seen in 80% of FP cells (25/31 cells from 17 embryos), showing that these structures are very common in FP cells but can be missed in Δt2min or Δt5min movies due to their short lifetime. Indeed, we found that digitations had a median lifetime of 50sec (FigS2b). In many cases, we could witness both digitation extension and retraction. Their median length at their maximal extension was 2µm (Fig. S2b) and almost all digitations pointed towards the BB (95%, 90/95 digitations, 25 cells from 14 embryos) and about 45% of them touched the BB. As previously mentioned, almost all digitations extended from transverse membranes (92%, 88/95, FigS2a) and most digitations formed at a spot were we could previously see another digitation (“recurrent” digitations, 85%, 44/52); the median time lapse between two successive digitations was 70sec (Fig. S2b). As with our Δt2min and Δt5min movies, there was not a good correlation between digitation position (anterior or posterior membrane) and BB movements: posterior digitations were followed by a posteriorward BB movement in 40% of cases (16/40) whereas anterior digitations were followed by an anteriorward BB movement in only 30% of cases (13/34) (Fig. S2c) suggesting that these digitations are not responsible for BB movements but rather a consequence of the forces exerted on BBs.

### Dynamic microtubules link BBs and transverse membranes

Since centrosomes are the main microtubule organizing centers of animal cells and their positioning in many systems has been shown to depend on microtubules we then set out to investigate microtubule dynamics within the apical surface of FP cells in early stage embryos. Live-imaging of microtubules with EB3-GFP revealed a highly dynamic network of microtubules originating from the centrosome/BB and directed to apical junctions (FigS2d, Movie S7). The time interval between an EB3 comet coming from the BB touching a spot at the transverse membrane and the occurrence of either a digitation at this spot or a BB movement towards it was very short: 10 sec in 50% of the cases and less than 1 min in 95% of the cases (Fig. S2e). These results show the existence of dynamic microtubules linking the BB and particular spots of the apical transverse membranes before BB movement toward these spots and/or digitation formation at these spots. This suggests that the mechanical forces responsible for the back-and-forth BB movements between the anterior and posterior transverse membranes are mediated by dynamic microtubules.

Overall, our dynamic analysis reveals a highly motile behavior of BBs in FP cells at early somite stages. This was unexpected, given that FP cells are already ciliated at these stages (Fig. 1b) (^5^). As somitogenesis proceeds, BB motility decreases. BBs progressively stop shuttling from anterior to posterior cell junctions and their contacts with the posterior membrane last longer. We also uncover membrane digitations forming at precise spots of transverse apical membranes and show that these spots are linked to BBs by dynamic microtubules. These results suggest the existence of a molecular complex at precise spots of transverse membranes, probably at the level of apical junctions, which is able to exert mechanical forces on the BB via microtubules, and whose distribution becomes biased to the posterior side of each FP cell as development proceeds.

### Posterior enrichment of Par3 precedes BB/posterior membrane contact

In Drosophila, the apical junction protein Par3/Bazooka modulates centrosome positioning in the male germline and embryonic ectoderm (^24, 25^). In order to test a potential role for Par3 in BB posterior positioning in FP cells, we first assessed Par3 localization by immunostaining (Fig. 4a). At the 14 s stage, Par3 localized at apical junctions of FP cells (Fig. 4a). Par3 patches were detected on transverse membranes and in close contact with posteriorly docked BBs (white arrows, Fig. 4a). This distribution was confirmed using the BazP1085 antibody (Fig. S3a), which recognizes a conserved Par3 phosphorylation site (^26^). Par3 patches were also present in FP cells in which the BB was not yet in contact with the posterior membrane (Fig. 4a and S3a right panels) showing that this enrichment precedes stable BB/posterior membrane contact.

**Figure 4.**
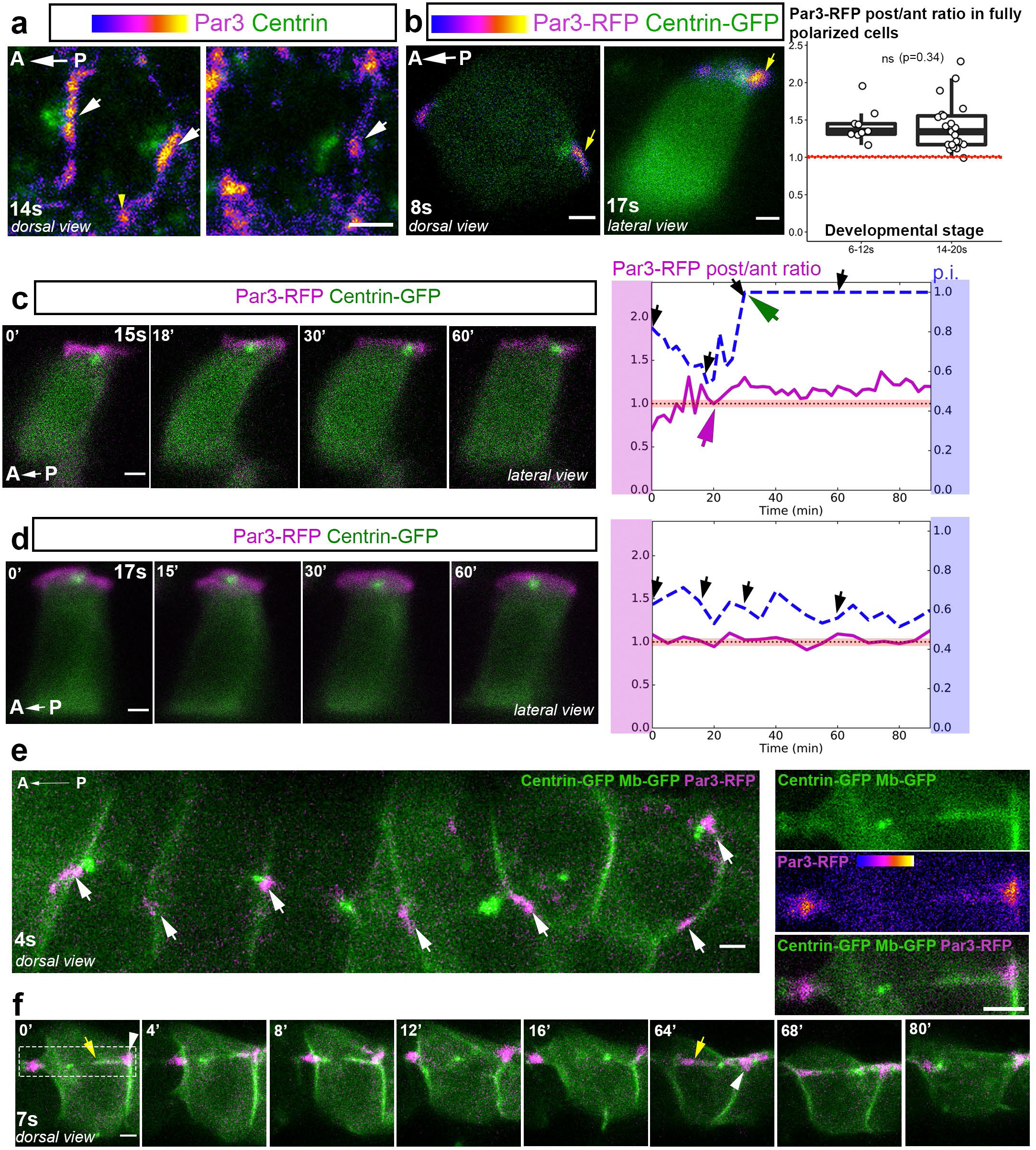
Par3 is asymmetrically localized in FP cells and forms patches at which almost all BB/membrane contacts occur. **a)** Individual cells from dorsal views of 14 s stage embryos showing IF with a Par3 antibody in FP cells. Two distinct cells are shown. Par3 localizes at apical junctions and is enriched at tricellular junctions (yellow arrowhead) and in patches at transverse membranes (white arrows), whether the BB is in contact with the posterior membrane (left images) or not (right image). **b)** Representative images of mosaically labelled FP cells expressing Par3-RFP and Centrin-GFP at early (8 s, left) or late (17 s, right) stages. Yellow arrows point at posterior Par3-RFP enrichment. Boxplots on the right show quantification of Par3-RFP posterior/anterior fluorescence intensity ratio in fully polarized FP cells at early and late stages. The red dotted line indicates a ratio of 1 (corresponding to a symmetric Par3-RFP distribution) (6-12s, mean ratio= 1.42, 7 embryos, 9 cells; 14-20s mean ratio =1.38, 13 embryos, 21 cells, Wilcoxon test). **c** -**d)** Images of time-lapse movies showing individual FP cells from embryos mosaically expressing Par3-RFP (magenta) and centrin-GFP (green) (lateral view). Par3-RFP posterior/anterior fluorescence intensity ratio is plotted on the right plots (magenta curve) along with the polarization index (« p.i. », dashed blue curve). Black arrows on plots indicate the time-points corresponding to the images displayed on the left. **c)** FP cell with Par3 posterior enrichment in an embryo filmed between the 15 s and 17 s stages. Par3 posterior enrichment starts 20 min after the beginning of the movie (magenta arrow), 10 min before BB/posterior membrane contact (green arrow). **d)** FP cell with no posterior Par3 enrichment (Par3-RFP post/ant ratio close to 1) with a BB oscillating around the middle of the apical surface, in an embryo filmed between 17 s and 19 s. **e-f)** Images from time lapse movies of early stage embryos mosaically injected with centrin-GFP (green), Membrane-GFP (green) and Par3-RFP (magenta) mRNAs. All pictures are dorsal views of FP cells. **e)** global view of 6 adjacent FP cells; white arrows point at Par3 patches (aligned along the AP axis) with which BB will make contacts during the movie. **f)** Example of a BB moving back and forth and contacting the membrane at Par3 patches. Posterior and anterior membrane digitations originating from Par3 patches and partially coated with Par3 can also be seen. Yellow arrows point to posterior (t=0’) and anterior (t=64’) digitations. White arrowheads point to Par3 patches. Par3 patch deformation can be seen at t=64’ and at t=0’ (images on the right show a close-up on the framed region at t=0’). Scale bars: 2µm.

In order to test whether Par3 is asymmetrically enriched in FP cells, we used a mosaic expression approach of Par3-RFP and centrin-GFP in live embryos. Quantification of Par3 expression showed that, among fully polarized (p.i. =1) individual Par3-RFP expressing FP cells, both at early (6-12s, Fig. 4b, left cell) and late (14-20s, Fig. 4b, right cell) stages, almost all cells had a Par3-RFP post/ant ratio greater than 1 (Fig. 4b right plot) (29/30 cells, 20 embryos; 6-12s, mean ratio= 1.42, 14-20s mean ratio =1.38). To determine whether Par3 posterior enrichment preceded BB/posterior membrane contact, we imaged BB movements and quantified Par3-RFP posterior/anterior ratio at each time-point; we found that Par3-RFP was enriched posteriorly before BB/posterior membrane contact (Fig. 4c, d) (12/14 cells, 12 embryos) (Movie S8). In contrast, BBs of FP cells with weak or no posterior Par3 enrichment remained unpolarized (either making no contact (2/5 cells, 5 embryos) or unstable contacts (3/5 cells, 5 embryos) with the posterior membrane (Fig. 4d and Movie S9).

Thus, Par3 forms patches at FP apical transverse membranes and BBs are posteriorly docked at these patches. In addition, Par3 is enriched posteriorly before BB/posterior membrane contact. Together, our data strongly suggest that Par3 is a key player in BB posterior positioning.

### At early stages, BBs contact transverse membranes exclusively at Par3 patches

During the second half of somitogenesis, Par3 formed a continuous belt at apical junctions of FP cells, although it was locally enriched, forming patches that associated with centrosomes, as described above. In contrast, at the 4 to 8 s stages, Par3 formed discrete patches at FP apical transverse membranes, but not at lateral membranes. These patches were roughly aligned with the AP axis of the embryo (Fig. 4e, white arrows). Strikingly, BBs made contacts with anterior and posterior transverse membranes (as described in Fig. 1) exclusively at these patches (58 cells from 18 embryos) as shown in Fig. 4f and Movies S10 and S11.

In 40% of these cells (23/58), the discrete Par3 patches stretched towards the BB (for example, Fig. 4f yellow arrows) and was actually covering a membrane digitation originating from either the posterior (Fig. 4f, t=0’, see also inset on the right) or the anterior membrane (Fig. 4f, t=64’) and extending towards the BB (Movie S11). Of the 39 digitations we saw, 92% were located at a Par3 patch (36/39 digitations from 23 cells and 14 embryos)(see Movie S11 at t=32min for a rare example of a digitation not originating at a Par3 patch). The presence of membrane digitations and their overlap with Par3 patches point to the existence of mechanical forces between BBs and membranes at Par3 patches and suggests that Par3 could be required for local force generation. Strinkingly, this role of Par3 could be more general than just BB antero-posterior contacts, since in dividing FP cells at early stages, after cytokinesis the centrosomes always rapidly (within 10min) moved back towards Par3 patches, next to the midbody (9/9 cells from 9 embryos, Fig. S3b and Movie S12).

### Par3 over-expression or aPKC-mediated clustering defects disrupt BB positioning

To test whether Par3 is required for posterior BB positioning in the FP, we first used loss-of-function approaches. MO-mediated knock-down of Par3ab (also known as Pard3 or ASIP) did not disrupt FP PCP (Fig. S4a), nor could we see a defect in a *MZpar3ab* mutant (^27^) (Fig. S4c). However, in both cases, Par3 patches could still be detected in the FP by immunostaining (Fig. S4b, d-f), suggesting that *par3ab* loss-of-function was compensated for by its paralogous genes (*par3aa*, *par3ba* or *par3bb*), which could also be detected by our Par3 antibodies thanks to the high conservation of the epitopes. In situ hybridization for all *par3* genes showed that *par3aa*, *ab* and *ba* were broadly expressed in the neural tube during somitogenesis (Fig. S5a).

Combining MOs against these three genes did not lead to FP polarization defects, but Par3 patches were still present and their prominence and number not affected, despite a significant loss in Par3 immunostaining signal (Fig. S5b-f): we could not test the effect of higher doses of MOs on FP polarization due to a developmental arrest before somitogenesis.

We thus turned to a mild over-expression approach to disrupt Par3 posterior enrichment and patch formation. Over-expressed Par3-RFP in the FP localized to apical junctions and did not disrupt apico-basal polarity (Fig.5a). Par3-RFP over-expression disrupted BB posterior positioning in the FP (median p.i. of 0.8 versus 1 in Par3-RFP negative cells, Fig. 5a,). Furthermore, isolated Par3-RFP negative cells and Par3-RFP negative cells adjacent to Par3-RFP over-expressing cells had similar polarization indexes, showing that Par3 overexpression effect is cell autonomous (147 isolated negative cells and 391 non-isolated negative cells from 20 embryos, Wilcoxon test p-value=0.19).

**Figure 5.**
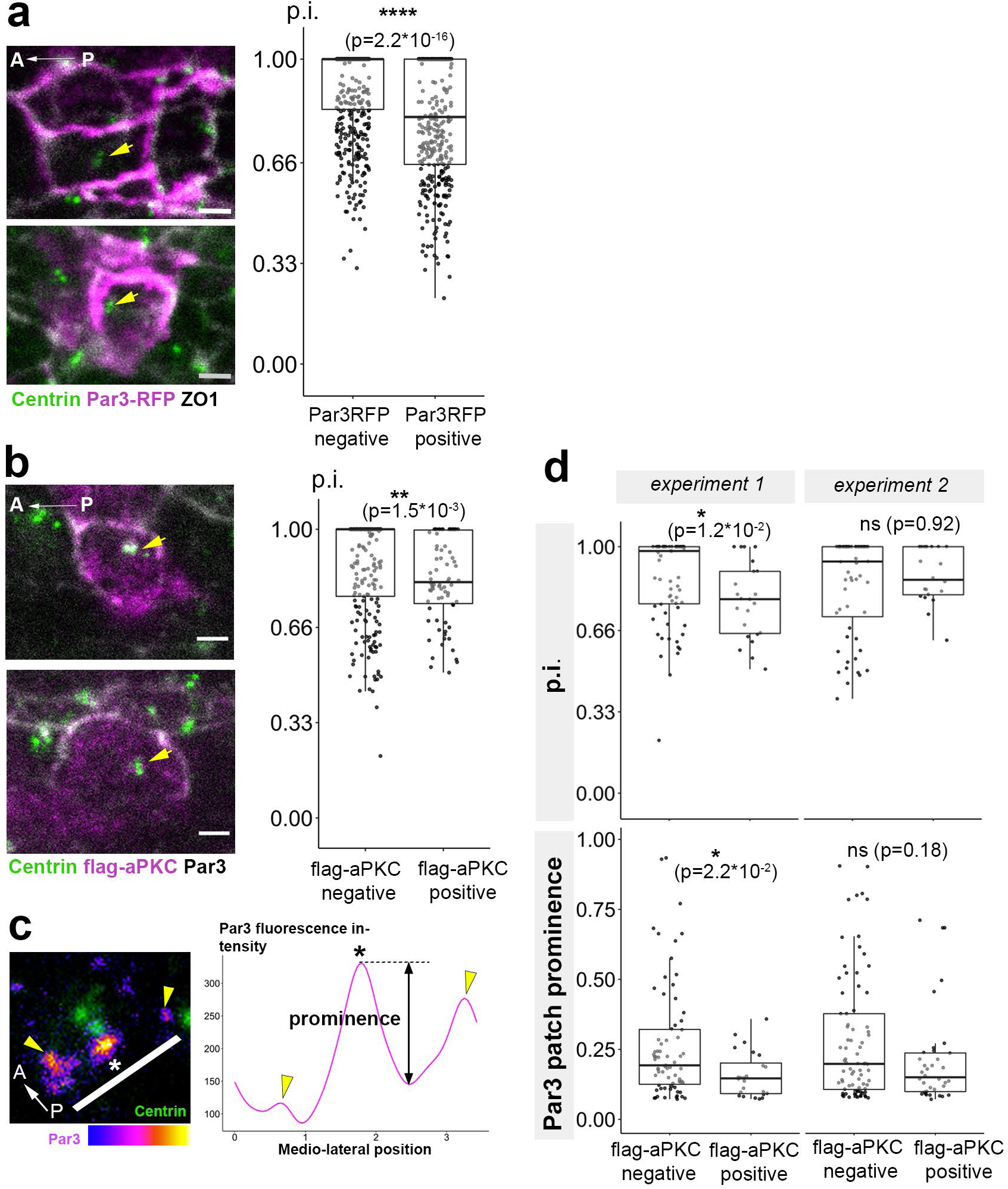
Disruption of FP polarization by Par3 or aPKC overexpression. **a)** Polarization index (p.i., cf Fig. 1) of Par3-RFP negative and positive FP cells from embryos mosaically over-expressing Par3-RFP. Representative immunostaining pictures of FP from embryos mosaically over-expressing Par3-RFP are displayed on the left; yellow arrows point at mispositioned BBs in Par3-RFP over-expressing cells. (538 Par3RFP negative cells and 375 Par3RFP positive cells from 20 embryos) **b)** Polarization index of flag-aPKC negative and positive FP cells in Netrin-KalTA4 embryos injected with a UAS:flag-aPKC construct (241 negative cells and 79 positive cells from 24 embryos). Representative immunostaining pictures of FP from embryos mosaically over-expressing flag-aPKC are displayed on the left; yellow arrows point at mispositioned BBs in flag-aPKC over-expressing cells. **c)** Par3 patches prominence is defined as the height of Par3 fluorescence peak relative to the highest and nearest valley (local fluorescence minimum). For each cell, prominence is normalized by the lowest Par3 intensity value. Right scheme: yellow arrows: tricellular junctions; white bar: orientation of the fluorescence measurement along the transverse membrane, star: position of Par3 patch **d)** Quantification of FP cell polarization index (upper plots) and Par3 patches prominence (bottom plots) in embryos mosaically expressing flag-aPKC. (experiment 1: 9 embryos, 23 flag-aPKC positive and 70 negative cells. experiment 2: 6 embryos, 20 flag-aPKC positive and 56 negative cells). Scale bar: 2µm. Comparison was done using a Wilcoxon test.

In order to confirm these results, we used another approach to disrupt Par3 endogenous distribution via aPKC over-expression (^25, 28^). We found that mosaically over-expressing aPKC with the KalTA4-UAS system (a Zebrafish-optimized GAL4-UAS system^29^) led to BB polarization defects similar to Par3 overexpression (Fig. 5b). In addition, in these experiments, the extent of Par3 localization defects correlated with that of BB polarization defects. In one experiment where we observed a significant decrease in Par3 patch prominence in aPKC overexpressing cells, we also observed a significant decrease in BB polarization (Fig. 5d, “experiment 1”). In another experiment where the change in Par3 patches prominence was present but not statistically significant, BB positioning was less affected (Fig. 5d, “experiment 2”).

Together, these results indicate that the extent to which the polarization is affected indeed depends on the strength of the effect on Par3 patches.

These results strongly suggest that Par3 posterior enrichment and patch formation are required for proper BB positioning in the FP.

### In the PCP mutant *vangl2* BB are still motile but make more contacts with lateral membranes

Vangl2, a core PCP protein, is involved in zebrafish FP PCP (^5^) but the downstream mechanisms linking Vangl2 to centrosome posterior positioning are unknown. We thus analyzed the dynamics of FP polarization in the *vangl2^m209^* mutant (^30^). At 18 s, the BB of *vangl2^m209/m209^* FP cells was mispositioned at the center of the apical surface, while *vangl2^m209/+^* and wt embryos had normally polarized BBs (median p.i.=0.6 versus 1 for wt or *vangl2^m209/+^*) (Fig. 7a, FP polarization plot). Live-imaging of *vangl2^m209/m209^* FP revealed several BB behavior changes in *vangl2*^m209/m209^.

At early stages in *vangl2*^m209/m209^ FP cells, BBs were still motile and even displayed an overall speed increase (Fig. 6d). Moreover, BB movements were still biased along the antero-posterior axis (Fig6b, 60% of total BB movements; 9 embryos, 20 cells, 220 movements), but BBs made more lateral movements and less antero-posterior movements compared to wt (10% less antero-posterior movements, Fig. 6a and 6b, Movie S13). The length of BB movements, which was smaller in the lateral than in the antero-posterior direction in wt, was equivalent in all directions in *vangl2*^m209/m209^ mutants (Fig. 6a).

**Figure 6.**
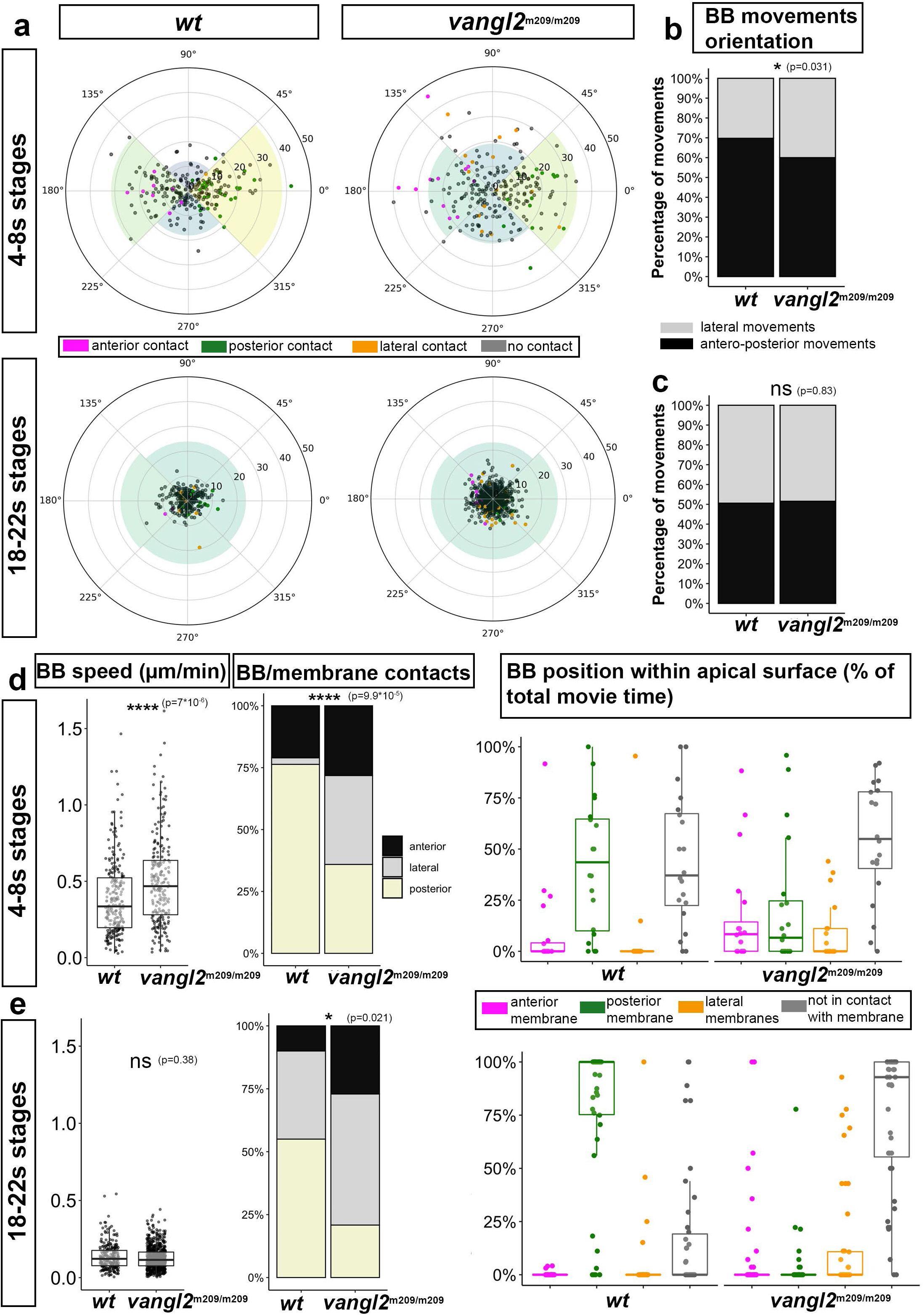
Abnormal BB behaviors in *vangl2^m209^* mutant FP. **a)** BB movements in wt (left column) and *vangl2^m209^* (right column) embryos at early (first line) and late (second line) developmental stages. Each dot represents the endpoint of a single BB movement, the starting point being the center of the circle; thus, the angles outside the circle represent BB movements orientation relative to the embryo anteroposterior axis, and the distance relative to the center of the circle represents its length (each circle radius corresponds to 7 µm). The color of the dots indicates whether a movements leads to a membrane contact and, if so, the nature of the contact (anterior, posterior or lateral). (early stages: wt 8 embryos, 20 cells, 238 movements; *vangl2^m209^*: 9 embryos, 20 cells, 220 movements. Late stages: wt 4 embryos, 19 cells, 255 movements; *vangl2^m209^*: 4 embryos, 42 cells, 708 movements). **b, c)** Orientation of BB movements in wt and *vangl2^m209^* embryos at early (b) or late (c) stages. **d, e)** BB movements speed, nature of membrane contacts and total time spent in contact with membranes at early (d) and late (e) stages. Statistical tests: Wilcoxon test for comparison of BB speeds; Fisher test for comparison of BB movements orientation and BB/membrane contacts.

Despite the preserved antero-posterior bias in BB movements, *vangl2* mutants showed a striking loss of antero-posterior bias in BB/membrane contacts. The overall proportion of BB movements resulting in BB/membrane contacts was the same as in wt (around 16% of total BB movements), but the positions of these contacts were very different: in wt, most contacts occurred with the posterior membrane (75%) and almost none with the lateral membranes (3%), whereas in *vangl2*^m209/m209^, BB contacts occurred equally with anterior, posterior or lateral membranes (around 33% each) (Fig. 6d, middle barplot). In addition, *vangl2*^m209/m209^ BBs spent less of their time in contact with either anterior, posterior and lateral membranes (Fig. 6d, right plot).

At later stages, almost half of the BBs remained in contact with the posterior membrane in wt, as previously described (Fig. 2e) (posteriorly docked BBs). In *vangl2*^m209/m209^ embryos, the vast majority of BBs did not stably dock at any membrane (except for 2 BB out of 42, docked at the anterior membrane). Instead, as suggested by our immunostaining, BBs remained at the center of the apical surface.

The few BB movements of wt cells as well as the many BB movements seen in *vangl2*^m209/m209^ cells were much smaller than at early stages (which is illustrated by a decrease in BB speed (Fig. 6e)) and had no preferential orientation (Fig. 6a). BB/membrane contacts were half less frequent than at early stages, both in wt and *vangl2*^m209/m209^ (around 7% of total BB movements), but we could still detect a significant difference in their position between wt and *vangl2*^m209/m209^, with more lateral and anterior contacts in *vangl2*^m209/m209^ (Fig. 6e, middle barplot).

### BB behavior defects in *vangl2* mutants are associated with abnormal Par3 clustering and localization

Since in wt embryos BBs only made contacts at Par3 patches, we wondered if it would still be the case in *vangl2* mutants. Par3 localized at apical junctions in *vangl2^m209/m209^* as in wt embryos (Fig. 7b). However, quantification of Par3 patches along the transverse apical junctions revealed a significant difference in the number and prominence of these patches. In wt, 90% of FP cells had at least a major Par3 patch (Fig. 7b, yellow arrows), with 39% of cells also having smaller secondary patches (Fig. 7c), while in *vangl2^m209/m209^* embryos, the number of FP cells with at least one phospho-Par3 patch was unchanged (around 90% of cells) but the number of cells with more than one patch was increased (54% of cells). In addition, the prominence of phospho-Par3 patches fluorescence intensity was decreased in *vangl2^m209/m209^* embryos (see Fig. 5c for prominence definition and quantification). Thus, Par3 forms more numerous and smaller patches in *Vangl2* mutants, showing a role for Vangl2 in Par3 clustering.

**Figure 7.**
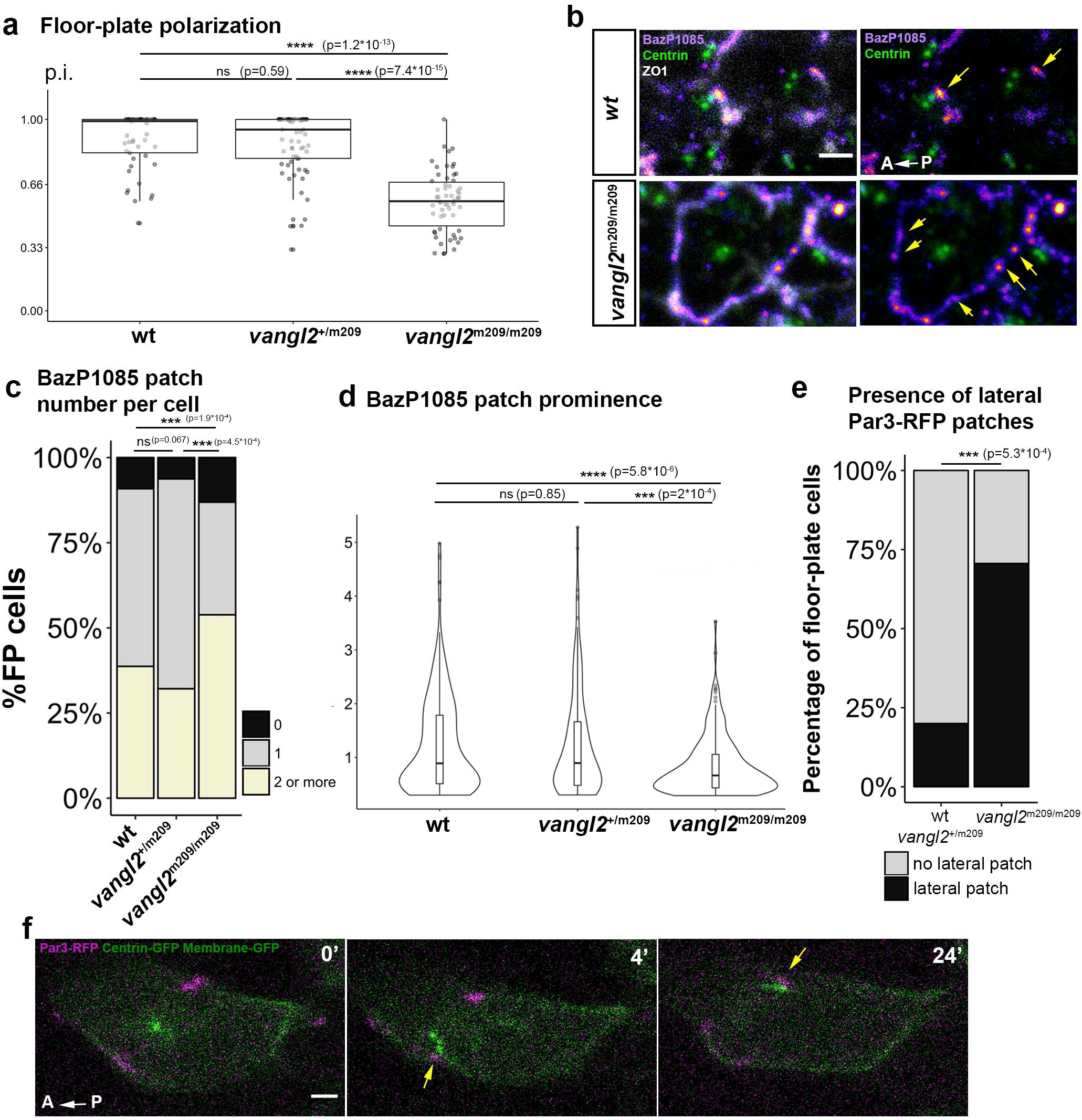
Par3 clustering and localization in *vangl2^m209^* mutant FP. **a)** Polarization index of *vangl2^m209/m209^* determined from immunostaining data wt: 2 embryos, 49 cells; *vangl2^m209/+^* : 3 embryos, 66 cells; *vangl2^m209/m209^*: 5 embryos, 57 cells. **b)** Immunostaining of phoshorylated Par3 (BazP1085 antibody) in *vangl2^+/+^* (wt) and *vangl2^m209/m209^* embryo FP at 18 s. In each case ZO1 staining was removed in the right image to reveal Par3 patches (yellow arrows). **c)** Quantification of the number of Par3 patches per cell on transverse membranes from immunostaining data as shown in b. d) Prominence of Par3 patches (BazP1085 antibody) in wt and *vangl2^m209/m209^* mutant embryo FP at 18 s In a-d, *vangl2^+/+^* : 7 embryos, 186 cells; *vangl2^m209/+^* : 5 embryos, 112 cells; *vangl2^m209/m209^* : 7 embryos, 129 cells. **e)** Percentage of cells displaying a lateral Par3-RFP patch in live-imaging experiments (such as the one described in f). *vangl2^+/+^* and *vangl2^m209/+^* : 16 embryos, 45 cells; *vangl2^m209/m209^* : 7 embryos, 17 cells. **f)** Images from movies of 5s *vangl2^m209/m209^* embryos mosaically injected with Par3-RFP, Centrin-GFP and Membrane-GFP mRNA at the 16-32 cell stage. Yellow arrows point at contact events between lateral Par3 patches and BBs. Statistical tests: Wilcoxon test for comparison of p.i. and prominence; Fisher test for comparison of patch number and percentage of cells with lateral patches.

To further analyze a potential link between abnormal BB behavior and Par3 patches mis-localization in *vangl2*^m209/m209^ embryos, we made time-lapse movies of mutant embryos mosaically injected with Par3-RFP (Fig. 7f) (Movie S13). In *vangl2* mutants, FP cell BBs contacted the membrane only at Par3 patches (Fig. 7e), as seen in wt, suggesting that Vangl2 did not directly affects the ability of Par3 patches to attract BBs. However the distribution of Par3 patches was very different. Early *vangl2^m209/m209^* embryos displayed many more cells with lateral Par3 patches compared to wt (70% vs 20%, Fig. 7g). In addition, they had more cells with an anterior Par3 patch (82% vs 67%) and less cells with a posterior patch (65% vs 87%).

These results strongly suggest that, in *vangl2*^m209/m209^ embryos, abnormal BB behavior and polarization failure are due to the fragmentation and mispositioning of Par3 patches along the apical junctions of FP cells.

## DISCUSSION

In this paper we have analyzed the dynamics of BB posterior positioning in the embryonic zebrafish FP. We show that, during early somitogenesis, BBs are highly motile and able to contact apical junctions several times per hour. As somitogenesis proceeds, BBs settle down posteriorly at junctions enriched in Par3, and we show that Par3 enrichment is essential for BB posterior localization. In the PCP mutant *Vangl2*, BBs show poorly oriented movements and this correlates with Par3 signal fragmentation and spreading to lateral junctions (Fig. 8a). Our data lead us to propose a model in which Par3 posterior enrichment, controlled by the core PCP pathway, increases microtubule-driven pulling forces from the posterior side, which eventually results in BB docking at the posterior Par3 patch (Fig.8b).

**Figure 8.**
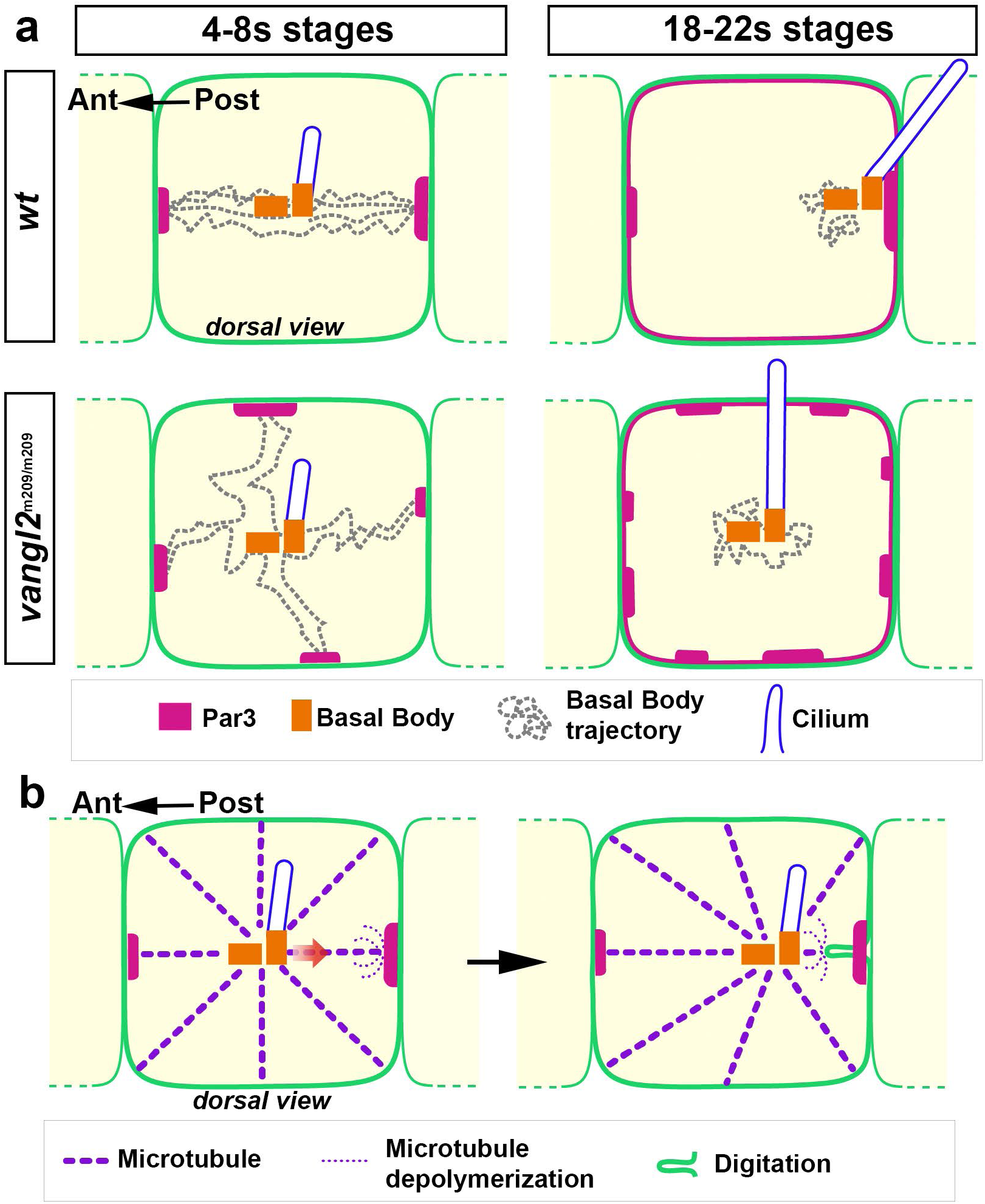
Summary of the main findings and hypothetical mechanism. **a)** Scheme showing BB behavior at early and late stages in the FP of wt and vangl2 mutants. In wt at early stages of polarization, the BB makes back and forth movements (dotted lines) between anterior and posterior Par3 patches (magenta) but very few lateral movements. In contrast, in *vangl2* mutants, BBs make more lateral movements and contacts and these contacts always occur at ectopic lateral Par3 patches. At later stages, the BB displays low motility. In wt, BBs are either docked to posterior Par3 patch or move in the posterior side of the cell, close to the patch, whereas in *vangl2* mutants BBs move around the center of the apical surface, which is probably due to earlier polarization defects and the still abnormal dispersion of Par3 patches around the apical junctions. Note that in both wt and *vangl2* embryos, at all stages studied, cilia are present and do not seem to impair BB movements. **b)** Hypothetical microtubule-based mechanism for BB polarization. Within the apical surface, dynamic microtubules emanate in all direction from the BB but can be captured at Par3 patches, at which their depolymerization is triggered (in this case at the posterior patch). The resulting mechanical force pulls on the membrane and the BB, which can result in digitation formation and/or BB movement towards the patch. The posterior Par3 enrichment makes it more likely for posterior pulling to happen (compared to anterior pulling) which ultimately results in a stably docked posterior BB at later stages.

### Ciliary basal bodies exhibit high motility in FP cells of wt and vangl2 mutant embryos

Analysis of fixed samples showed that posterior positioning of BBs within the apical surface of FP cells progressed regularly within the 8 hours time frame of our study and was complete at the 18 s stage. Surprisingly, live imaging revealed that BBs make fast back and forth movements within the apical surface. This high motility of the BB (median speed 0.2 µ/min at 4-8s) is unexpected given the presence of a growing cilium anchored to its distal part. It contrasts with the situation in the mouse cochlea, where live-imaging of explants have suggested very slow and regular movements of the BBs to the lateral cortex of inner hair cells (estimated speed of 10-50 nm/h) (^19^). BB motility decreases at later stages of polarization, coincident with their posterior docking. Live imaging also uncovered a clear antero-posterior bias in BB movements, indicating that the orientation of the forces underlying these movements is biased along the polarization axis from early stages on. However, we could not detect a significant global posteriorward bias in these movements in early-stages FP cells (Fig. 6a).

In fixed samples of *vangl2* mutants, the BBs remain at the center of the FP cell apical surfaces (^5^ and Fig. 7a-b), suggesting a decrease in their motility. Strikingly, our dynamic analysis revealed that in *vangl2^m209/m209^* embryos, BBs are still highly motile. Major differences with wt BBs are that they make many contacts with lateral membranes, whereas wt BBs do not, and that their contacts are shorter than for wt BBs. This suggests that there are molecular cues at the membrane organizing/driving BB movements, and that these cues are still present but disorganized in space in PCP mutants.

Thus, our live imaging study uncovers a totally unexpected motile behavior of BBs during a cilia planar polarization process. It will be interesting to investigate whether this behavior is conserved in other tissues undergoing cilia translational polarity.

### Par3 cortical patches recruit the BB

We proposed the asymmetric maturation of cell junctions as a possible cause for posterior BB positioning. Accordingly, we found that Par3 accumulated in patches at the posterior apical junctions of FP cells before BB posterior docking. Moreover, we showed that perturbed Par3 localization affected BB polarization. Interestingly, in the Drosophila early gastrula ectoderm, Par3 isotropic distribution around apical junctions contributes to epithelial integrity, but in aPKC loss of function mutants, Par3 accumulates as discrete patches that align along the dorso-ventral axis and “recruit” centrosomes (^25^). Centrosome docking at discrete Par3 patches has also been observed in Drosophila germ stem cells and is critical for proper division orientation (^24^). Together with these published data, our results on BB positioning in zebrafish FP strongly suggest that Par3 may be broadly involved in recruitment of centrosomes and BBs in different systems.

Our live imaging data strongly suggest that Par3 is involved in generating mechanical forces on BBs to pull it toward the membrane. First, the BB contacts the membrane exclusively at Par3 patches. Second, membrane deformations support the existence of mechanical forces between Par3 patches and BBs. Third, the predominance of posterior digitations over anterior ones (Fig. 3b) suggests that more force is exerted on the BB from the posterior side, where Par3 is enriched. Such membrane digitations have been previously observed during cell division in the *C. elegans* zygote (^31^), in the *C. intestinalis* embryo ectoderm (^32^) and in rare cases at the immunological synapse (^33^). In all cases, the existence of pulling forces between the centriole and the membrane has been proposed.

We propose that digitations are a consequence of mechanical forces between the BB and Par3 patches rather than a driver of BB movement (as was previously hypothesized in Ciona embryos (^32^)). First, digitations were rare (even in Δt10sec movies, only half of BB movements were associated with digitations) and second, there was not a good correlation between the location of a digitation and the direction of BB movement. One can wonder why digitation formation occurs for some BB movements and not others. A possibility is that digitation formation depends both on mechanical forces pulling the membrane and on cortical stiffness. BB movement and digitation formation could thus depend on the balance between these two forces. It would therefore be interesting to investigate the dynamics of cortical actin during BB movements and digitation formation.

### Possible role of microtubules in BB recruitment to Par3 patches

Our results suggest that mechanical forces between Par3 and the BB could be exerted by microtubules. Dynamic microtubules link the BB and Par3 patches before BB movements and/or digitation formation. Microtubules are required for membrane digitations formation in several systems (^31, 32^) and are thus very likely to transmit the mechanical forces between BB and Par3 patches that lead to BB movements and/or membrane digitations.

An interesting further question concerns the mechanisms that regulate microtubule dynamics to lead to BB movements. BB movements towards Par3 patches could involve local microtubule depolymerization at the patch (Fig. 8b), coupled to microtubule anchoring by dynein as proposed for the migration of the centrosome toward the immunological synapse (“end-on-capture-shrinkage” mechanism)(^33^). Indeed, Par3 can interact with Dynein (^34^) and also with microtubules, directly (^35^) or indirectly via 14-3-3 proteins (^36^). Consistent with a role for cortical dynein, a recent study in mouse ependymal multi-ciliated cells showed a role of cortical dynein in the off-centering of BB clusters (^37^). Moreover, dynein cortical localization depends on Daple, which is a known partner of Par3. Par3 could also regulate microtubule depolymerization via Rac1, which mediates Par3 function in the mouse cochlea (^38^). In different systems, Par3 regulates the local activity of Rac via the RacGEFs Tiam1 and Trio (^39,40,41^). Par3 can increase microtubule catastrophe rate by inhibiting Trio in neural crest cells (^42^), and Rac1 can regulate microtubule dynamics via CLIP-170 or Stathmin in other systems (^43, 44^).

Interestingly, microtubules also actively maintain BB polarity at later stages, as recently demonstrated (^21^), but the authors could not distinguish between a role of microtubules as mechanical forces generators or as tracks for PCP proteins transport and asymmetric localization: indeed in nocodazole-treated embryos, Vangl2 asymmetric localization is lost. This is an important caveat for studies that will try to address the role of microtubules in BB positioning in a PCP context in more details, since microtubules role in PCP protein transport and asymmetric localization seems to be widely conserved (^45, 46^). Finally, Par3 apical localization also depends on microtubules in zebrafish embryo neural tube (^47^): thus it might prove difficult to disentangle the different potential roles of microtubules in the asymmetric positioning of BBs of planar polarized ciliated epithelia.

### The core PCP protein Vangl2 is involved in BB positioning via Par3 enrichment to the posterior membrane

In *vangl2^m209/m209^* embryos, BBs show behavioral defects, making more lateral movements and more contacts with lateral membranes than in wt embryos. Strikingly, in *vangl2* mutants as in wt, BBs always contacted the apical junctions at Par3-positive patches. The altered behavior of BBs in *vangl2^m209/m209^* embryos correlated with a mislocalization of Par3 around the apical junctions of FP cells. Since Par3 mislocalization affected BB polarization, we propose that Par3 posterior enrichment and patch formation under the control of the PCP pathway is a main actor in BB posterior positioning (Fig.8a).

How PCP proteins act on Par3 localization in the FP remains to be uncovered. In FP cells, Vangl2 localizes anteriorly (^48^); thus, Vangl2 effect on Par3 could be mediated by Dvl. Indeed, Vangl2 is required for proper asymmetric localization of Dvl in planar polarized tissues and Dvl can recruit Par3 to the posterior membrane in Drosophila sensory organ precursors (^49^). Dvl could also recruit Par3 via Daple, as this protein colocalizes with Par3 in the mouse cochlea and can bind both Dvl and Par3 in yeast two-hybrid assays (^50^). Recent studies have shown that Par3 is asymmetrically localized within the plane of the epithelium in several systems, like Drosophila ommatidia (^51^) and sensory organ precursors just before asymmetric division (^52^), in Xenopus embryo ectoderm (^53^) and in the mouse cochlea (^38^), suggesting that in addition to its classical role in apico-basal polarization, Par3 might also be involved in PCP across species and even considered a bona fide core PCP protein. It will be interesting to investigate whether this Par3 posterior enrichment involves a positive feedback loop between BB and Par3 patches either directly by contact (^54^) or via dynamic microtubule (+) ends (^55^).

Finally, as asymmetric centriole positioning is now recognized as a conserved readout of PCP (^56, 57^), it will be interesting to investigate whether Par3 has a conserved role in centriole/BB positioning in other species where BB/centriole off-centering has been described and also depends on PCP proteins, for example in the embryo of the jellyfish Clytia *hemisphaerica (*^58^) or in Drosophila pupal wing (^56^).

## MATERIALS AND METHODS

### Zebrafish handling and experimentation

Wild-type and mutant zebrafish embryos were obtained by natural spawning. We used wild-type AB or (TL x AB) hybrid strains, *vangl2*^m209^ mutants (^30^, ZDB-GENO-190204-5), *par3ab fh305* mutants (^27^, ZDB-FISH-150901-20689) and created a transgenic Netrin-KalTA4 line, expressing the KalTA4 transcriptional activator in floor-plate cells. The Netrin-KalTA4 line was generated by injecting at the 1 cell stage 15pg of pNetrin-KalTA4 plasmid along with 20pg of Tol2 mRNA. To obtain the early stages (4-8s), embryos were collected at 10 am and incubated for 9 h in a 33°C incubator. To obtain later stages (14-20s), embryos were collected at 10 am and incubated for 2 h at 28 °C before being placed overnight in a 24 °C incubator. All our experiments were made in agreement with the European Directive 210/63/EU on the protection of animals used for scientific purposes, and the French application decree ‘Décret 2013-118’. The projects of our group have been approved by our local ethical committee ‘Comité d’éthique Charles Darwin’. The authorization number is 2015051912122771 v7 (APAFIS#957). The fish facility has been approved by the French ‘Service for animal protection and health’ with approval number A-75-05-25.

### Plasmid construction

**pUAS:flag-aPKC**: the rat aPKC (PKC zeta) coding sequence was amplified by PCR from addgene plasmid #10799 using primers (flagaPKC-forward and reverse) with 15bp 5’ and 3’ overhangs. Primers homologous to these overhangs (UASRho-forward and reverse) were used to amplify the backbone of the pUAS-RhoAwt plasmid (Hanovice 2016) (thus removing the mCherry-RhoA sequence). The two DNA fragments were then ligated into one plasmid with the InFusion HD cloning kit (Takara).

**pNetrin:Kal TA4**: a 1.4kb fragment from pCSKalTA4 comprising the KalTA4 promoter was amplified using the KalTA4-forward and KalTA4-reverse primers, then digested with XhoI and NotI (NEB) and placed under the control of the floor-plate specific 898bp Netrin1 enhancer (^59^) from a XhoI/NotI-digested pNetrin898:membCherry plasmid (Marie Breau, unpublished) by ligation with the T4 DNA ligase (NEB).

### mRNA, morpholino and plasmids injection

mRNAs were synthesized from linearized pCS2 vectors using the mMESSAGE mMACHINE SP6 transcription kit (Ambion). The following amounts of mRNA were injected into one-cell stage embryos: 22 pg for Centrin-GFP, 40 pg for mbCherry (membrane Cherry) or Membrane-GFP (Gap43-GFP). For Par3-RFP mosaic expression, mRNAs were injected at the 16-cell stage in a single blastomere, using 50 pg for Par3-RFP live-imaging or 150pg Par3-RFP for over-expression experiments along with Centrin-GFP and membrane-GFP mRNAs at the same concentration as for one-cell stage injections. Par3ab-MO was injected at a concentration of 0.3 mM at one-cell stage. For triple MO injections (Par3aa, ab and ba coinjection) each MO was diluted to 0.25 mM. MO sequences are given in the supplementary methods.

For flag-aPKC mosaic experiments, 25pg of pUAS:flag-aPKC were injected in NetKalTA4 embryos at the 1 cell stage. This leads to mosaic expression due both to the stochastic expression of KalTA4 and the mosaic delivery of plasmid to some cells and not others.

### Immunostaining

For immunostaining, embryos were fixed in Dent fixative (80% Methanol, 20% DMSO) at 25°C for 2 h, blocked in 5% goat serum, 1% bovine serum albumin and 0.3% triton in PBS for 1 h at room temperature and incubated overnight at 4 °C with primary antibodies and 2 h at room temperature with secondary antibodies. The yolk was then removed and the embryo mounted dorsal side up in Vectashield medium on a slide. Imaging was done using a Leica TCS SP5 AOBS upright confocal microscope using a 63X oil lens. A list of antibodies is given in the supplementary methods.

### Live imaging

Embryos were dechorionated manually and mounted in 0.5% low-melting agarose in E3 medium. Movies were recorded at the temperature of the imaging facility room (22 °C) on a Leica TCS SP5 AOBS upright confocal microscope using a 63X (NA 0.9) water immersion lens. The embryos were mounted either with dorsal side up (for early stages or after 18s, when FP cells apical surface is quite large) or on the side (for most 13-18s embryos, when the apical surface of FP cells is narrower and when the images are more blurred when taken from a dorsal view, maybe because of the thickness of the overlying neural tube). The anterior side of the embryos was positioned on the left and their antero-posterior axis aligned horizontally. A z-stack was acquired every 2 or 5min (or in some rarer cases every 4min) for most analysis (Δt2min and Δt5min movies) and every 10sec for some movies (digitations analysis and microtubule dynamics in Fig.S2, Δt10sec) with a z-step of 0.3µm. For each time-point the z-stack extended from the most dorsal side of the notochord to neural cells above the FP for Δt2min and Δt5min movies but was narrower for Δt10sec movies (to allow fast acquisition and reduce photobleaching and photodamage). For embryos mounted on the side, the z-stack extended through all the width of the FP. In every case, the z-stack encompassed FP cells apical surface with the moving BBs. For each time-point we then make a z-projection from a 3µm thick substack that encompass the apical centrioles/BB.

### In situ hybridization

Embryos at the 16 s and 24 hpf were processed as previously described (^60^). Matrices for probe synthesis were synthesized by PCR using either adult genomic DNA (for *par3aa*, *ba* and *bb*) or cDNA from 18 s embryos (for *par3ab*). Primer sequences are available in the supplementary methods.

### Quantification and statistical analysis

All bar-plots, boxplot and violin plots and statistical tests (Wilcoxon or Fisher tests, see figure legends) were generated with R (version 3.3.2) and Rstudio (Version 1.1.463). ns (non-significant): p>0.05, *: p<0.05, **: p<0.01, ***: p<0.001, ****: p<0.0001.

### BB position and movements

In all our images, the antero-posterior axis (easily visualized thanks to the underlying notochord, whose cells have an elongated shape orthogonal to the antero-posterior axis) is horizontal and the anterior side of the embryo is toward the left. To assess the polarization of FP cells, we used FIJI to first make a z-projection of thickness 3µm around the centrioles. We then manually measured the distance “a” by drawing a line between the most posterior centriole and the posterior membrane that was parallel to the antero-posterior axis of the embryo (ie parallel to the horizontal axis of the image) and the distance “b” by drawing a similar line, at the same level and also parallel to the antero-posterior axis, between the anterior and posterior membrane (Fig. 1a, dorsal view). We used a similar method for embryos mounted on the side (Fig. 1a lateral view). The polarization index (p.i.) was then calculated as 1-(a/b). In some rare cases, the transverse membranes in our z-projection around the centrioles are blurry (Fig.2b); in these cases however, going slightly more basally through the z-stack allows us to see a sharper membrane which is usually located just beneath the boundary of the above blurry region.

To follow the evolution of polarization index, the distances “a” and “b” were measured manually at each time-frame in FIJI. These distances and the polarization index were then plotted using python matplotlib (Python 2.7.13) and analyzed with a custom python script to extract relevant information such as the frequency of contact with posterior membrane or percentage of total time spent in contact with posterior membrane. (Fig2 and FigS1).

For automatic tracking of BB movements (Fig5), BB were tracked using Image J TrackMate (^61^). The movements were then manually curated to only keep active BB movements and not BB movements due to global shift of cells (especially at early stages when convergence-extension movements are still important), and to indicate whether a contact with an anterior, posterior or lateral membrane occurred. The results were then processed using a custom python script to calculate each movement length and angle relative to the horizontal axis (ie the antero-posterior axis of the embryo) and plotted using Python Matplotlib and R ggplot2. We defined antero-posterior movements as the ones with angles relative to horizontal axis inferior to 45° (ie in the intervals [315-45°[and [135-225°[in Fig. 6a).

For all analyzes, lateral membranes are defined as the ones more parallel to the horizontal axis and transverse membranes as the ones more orthogonal to the horizontal axis. The transition between lateral and transverse (anterior and posterior) membranes is evidenced by “Y”-shaped tri-cellular junctions (as in Fig. 4a) or sharp turns in the membrane (as in Fig. 7f).

### Par3-RFP posterior/anterior ratio

Fluorescence intensity was measured along the anterior-posterior length of isolated labelled FP cells in FIJI. A custom python script was then used to extract the first quarter (cell anterior side) and last quarter (cell posterior side) of fluorescence intensity values, to determine the area under each curve (corresponding to fluorescence intensity), calculate the post/ant ratio and plot it along with the polarization index (see BB movements analysis section).

### Par3 peaks quantification

Fluorescence intensity from immunostained embryos was measured along FP cells transverse membranes and exported to Matlab (R2018a). For each cell, the “findpeaks” function was used to detect Par3 fluoresence peaks and measure their prominence, which was then normalized by the minimal Par3 fluorescence value along the junction.

### Additional softwares

Adobe Photoshop was used to assemble the figures. Fig. 1a was done in Microsoft Powerpoint and Fig. 8 with Adobe Illustrator.

## ACKNOWLEDGEMENTS

We are grateful to the aquatic animal and cell imaging facilities of the IBPS (Institut de Biologie Paris-Seine FR3631, Sorbonne Université, CNRS, Paris, France) for their technical assistance. We thank Teresa Ferraro for sharing her expertise in image analysis, Marie Breau for her help in setting up the live imaging protocol, Isabelle Anselme for participation in genotyping and Sophie Gournet for her help in the design of Fig. 8. We thank Paula Alexandre for the kind gift of Par3-RFP construct, Andreas Wodarz for the BazP1085 antibody, Maximilien Furthauer for the *vangl2^m209^* line. We thank Nicolas David and Marie Breau for critical reading and insightful comments on the manuscript. This work was supported by funding from the Agence Nationale pour la Recherche (ANR, project CILIAINTHEBRAIN to SSM) and the Fondation pour la Recherche Médicale (Equipe FRM DEQ20140329544 and EQU201903007943 funding to SSM). A.D was supported by fellowships from the Ecole Normale Supérieure de Cachan and from the Fondation ARC contre le Cancer. The authors declare no competing financial interests.

## SUPPLEMENTARY INFORMATION

### Methods

#### Antibodies

| Antibody | Source | Identifier | Dilution |
| --- | --- | --- | --- |
| Mouse monoclonal IgG2a<br>anti-centrin (clone 20H5) | Merck Millipore | # 04-1624<br>RRID: AB_10563501 | 1:200 |
| Mouse monoclonal IgG1<br>anti-ZO1 (clone ZO1-1A12) | Invitrogen | RRID: AB_2533147 | 1:400 |
| Mouse monoclonal IgG2b<br>anti-acetylated-tubulin<br>(clone 6-11B-1) | Sigma-Aldrich | #T 6793 RRID:<br>AB_477585 | 1:400 |
| Rabbit polyclonal anti-Par3 | Merck Millipore | #07-330<br>RRID:AB_11213581 | 1:200 |
| Rabbit polyclonal anti-<br>phosphorylated-Ser1085-Bazooka | <sup>26</sup> | N/A | 1:200 |
| Rabbit polyclonal anti-DsRed | Takara | # 632496<br>RRID:AB_10013483 | 1:400 |
| Mouse IgG1 anti-FLAG<br>(M2) | Sigma | F3165 | 1:100 |
| Goat anti-mouse IgG1<br>Alexa633 | Molecular probes | # A-21126<br>RRID:AB_2535768 | 1:400 |
| Goat anti-mouse IgG2a<br>Alexa568 | Molecular probes | # A-21134<br>RRID:AB_2535773 | 1:400 |
| Goat anti-mouse IgG2a<br>Alexa488 | Molecular probes | # A-21131<br>RRID:AB_141618 | 1:400 |
| Goat anti-mouse IgG2b<br>Alexa633 | Molecular probes | # A-21146<br>RRID:AB_2535782 | 1:400 |
| Goat anti-rabbit IgG<br>Alexa568 | Molecular probes | # A-11011<br>RRID:AB_143157 | 1:400 |

#### Oligonucleotides, Plasmides and Morpholinos

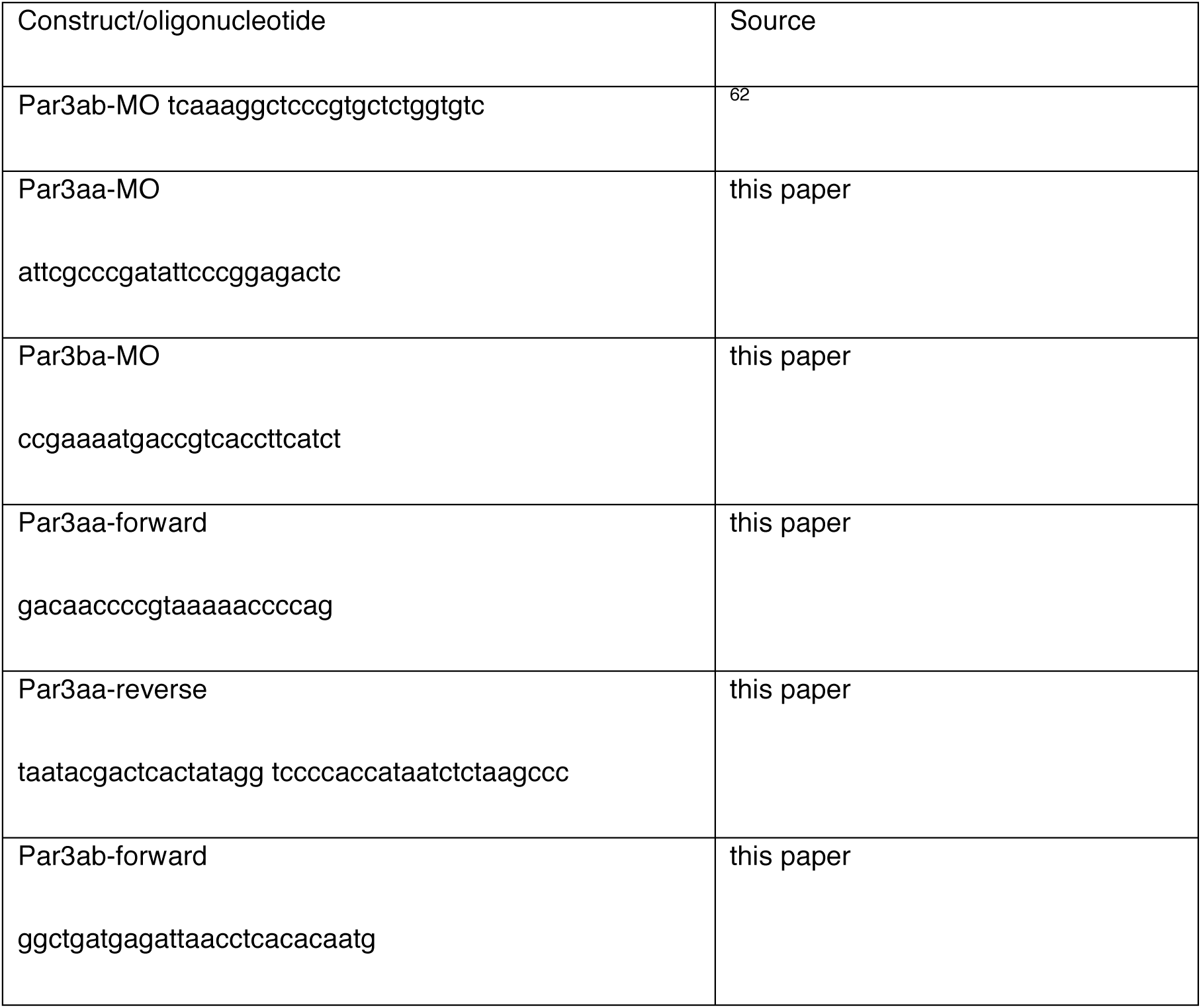
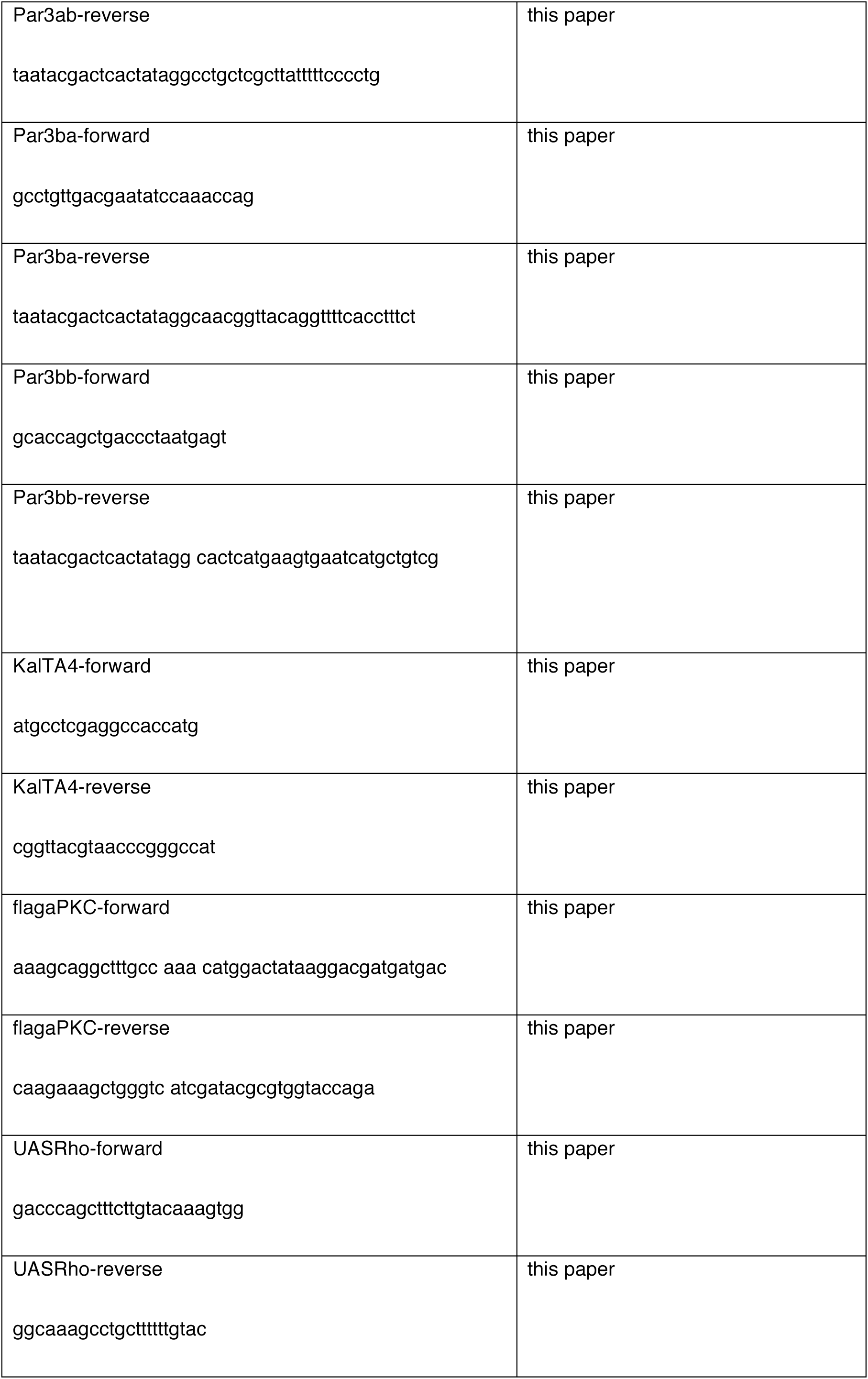
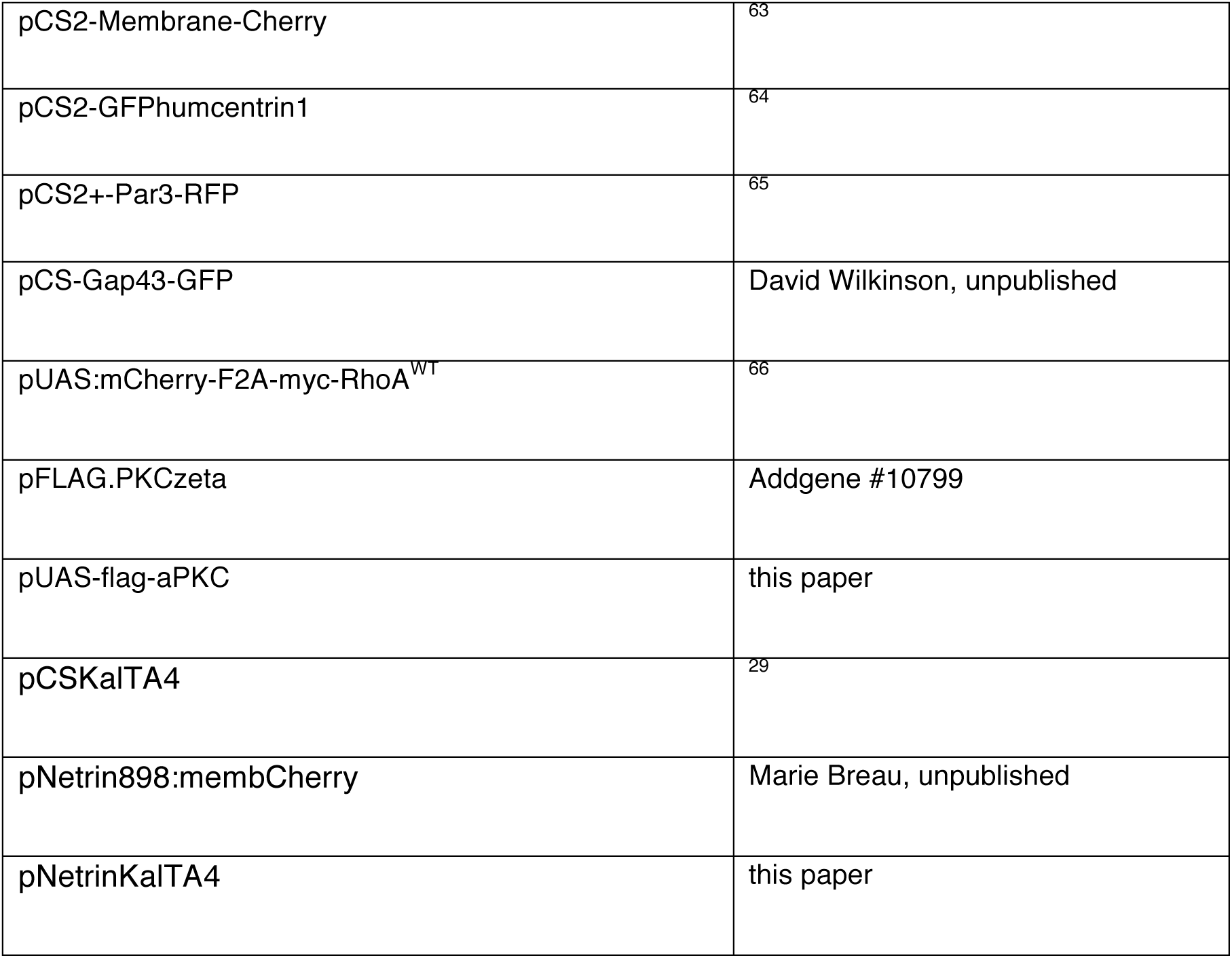

## SUPPLEMENTARY FIGURE LEGENDS

**Supplementary Figure S1:**
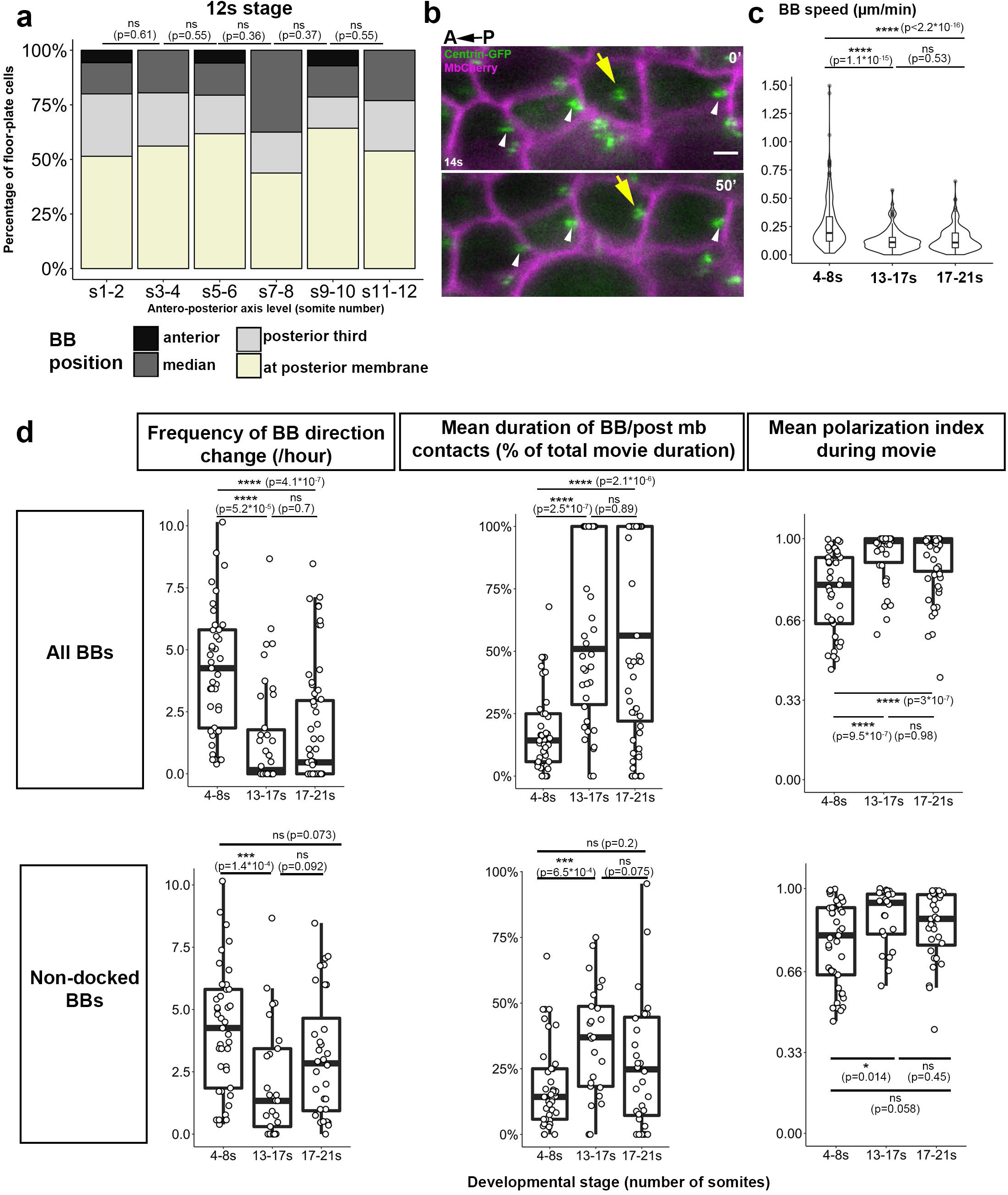
Further characterization of FP polarization in space and time. **a)** Quantification of FP polarization along the AP axis at 12 s. Analysis was performed on fixed immunostained embryos as described in Fig. 1a. Comparison between stages was done using a Wilcoxon test. **b)** Still images from FP BB (green) and membrane (magenta) live imaging (dorsal view, start at 14s stage). The yellow arrow points to BB that will move and make contacts with the posterior membrane between 0 and 50 min after the movie started. White arrowheads point at BBs in adjacent cells that stay in contact with the posterior membrane during this time interval. **c)** BB speed measured from live-imaging data at different developmental stages. The speed of each BB movement was calculated by dividing the value of BB/posterior membrane variations (corresponding to green curves in Fig. 1c-f) by the total duration of the movement (4-8s: 4 embryos, 38 cells; 13-17s: 6 embryos, 22 cells; 17-21s: 7 embryos, 32 cells). Comparison between stages was done using a Wilcoxon test. **d)** Movies described in Fig. 1 were used to quantify BB direction change frequency, mean duration of BB/posterior membrane contact events as a percentage of total imaging duration and mean polarization index during live-imaging. Plots in the first line take into account the BBs that stay in contact with the posterior membrane 100% of movie duration (posteriorly docked BBs) whereas the second line only represents BBs that are not posteriorly docked. Comparison between stages was done using a Wilcoxon test. Scale bar: 2µm.

**Supplementary Figure S2:**
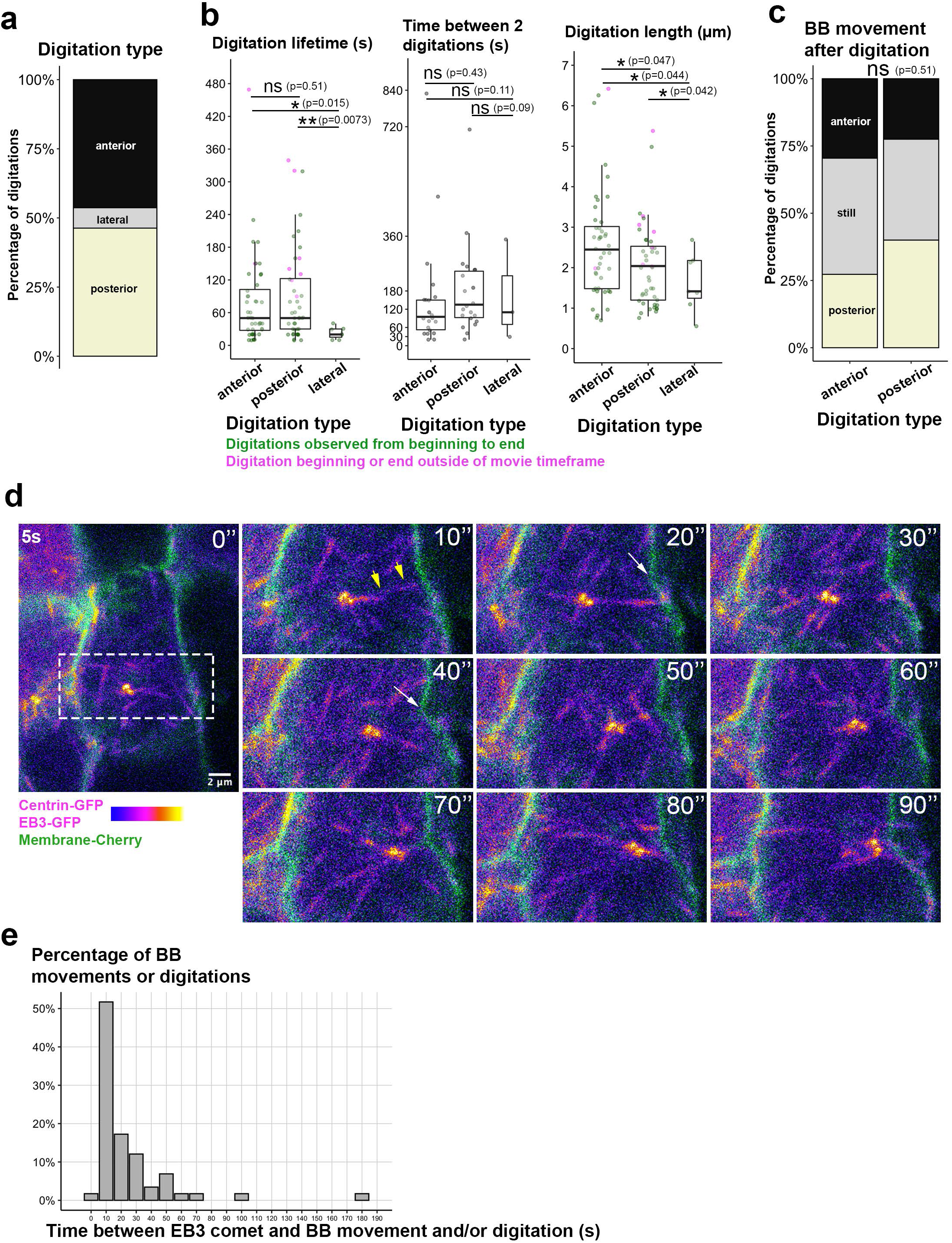
High temporal resolution imaging reveals numerous anterior and posterior digitations and dynamic microtubules between BB and target membrane. a-c) Characterization of digitations with high temporal resolution live-imaging (one image every 10sec) (12 embryos, 20 cells, 44 anterior digitations, 44 posterior digitations, 7 lateral digitations). **a)** Repartition of digitation types. Anterior, lateral and posterior correspond to the membrane from which the digitation forms. There were as many anterior as posterior digitations, in contrast to what we observe in our Δt=2 or 5 min movies (Fig. 3) **b)** Quantification of digitation lifetime, recurrence time (time elapsed before a particular digitation reforms at the same spot) and maximal length. Dots in magenta correspond to digitations that had formed before the beginning of live-imaging or had not disappeared yet when we stopped filming, and for which we could not determine the exact lifetime (which is thus under-estimated) (comparison done with Wilcoxon test). **c)** Direction of BB movements after an anterior (left) or posterior (right) digitation disappears. The BB either moved anteriorly, posteriorly or remained still (comparison done with Fisher test). There was no difference in digitation lifetime, which median value was 50sec both for anterior and posterior digitations. The time elapsed between two recurrent digitations was also not significantly different between anterior (median of 70sec) and posterior (median of 80sec) digitations. The presence of more posterior digitations in Δt2min and Δt5min movies but not in Δt10sec movies could be due to the fact that we miss most very short-lived digitations in Δt2min or Δt5min movies and that the short duration of our Δt10sec movies (in average 20min long) prevented us to see some long-lived digitations from extension to retraction (pink dots). Anterior and posterior digitations had similar size of around 2µm, although anterior digitations were slightly longer (2.4µm), probably due to the fact that BBs even at these early stages have a posteriorly biased position (Fig. S1d Mean polarization index plots). Posterior digitations were followed by a posteriorward BB movement in 40% of cases (16/40) whereas anterior digitations were followed by an anteriorward BB movement in only 30% of cases (13/34) suggesting that digitation formation is probably not a cause but rather a consequence of BB movements. **d)** Dorsal view of a FP cell of a 5s stage embryo mosaically injected with Centrin-GFP (BB), EB3-GFP (microtubule plus ends) and Membrane-Cherry, which was then imaged every 10s. The dotted frame corresponds to the enlarged region on the right. Yellow arrows (10’’): microtubule extending from the BB toward the spot of the posterior membrane that the BB will reach at t=90’’. White arrows: membrane digitation forming from the spot targeted by the microtubule highlighted at t=10’’ and toward the BB. **e)** Time elapsed between an EB3 comet reaching a membrane (such as in a at t=10’’) and the moment when a digitation forms or the BB starts moving towards this membrane (6 embryos, 10 cells, 58 EB3 comets).

**Supplementary Figure S3:**
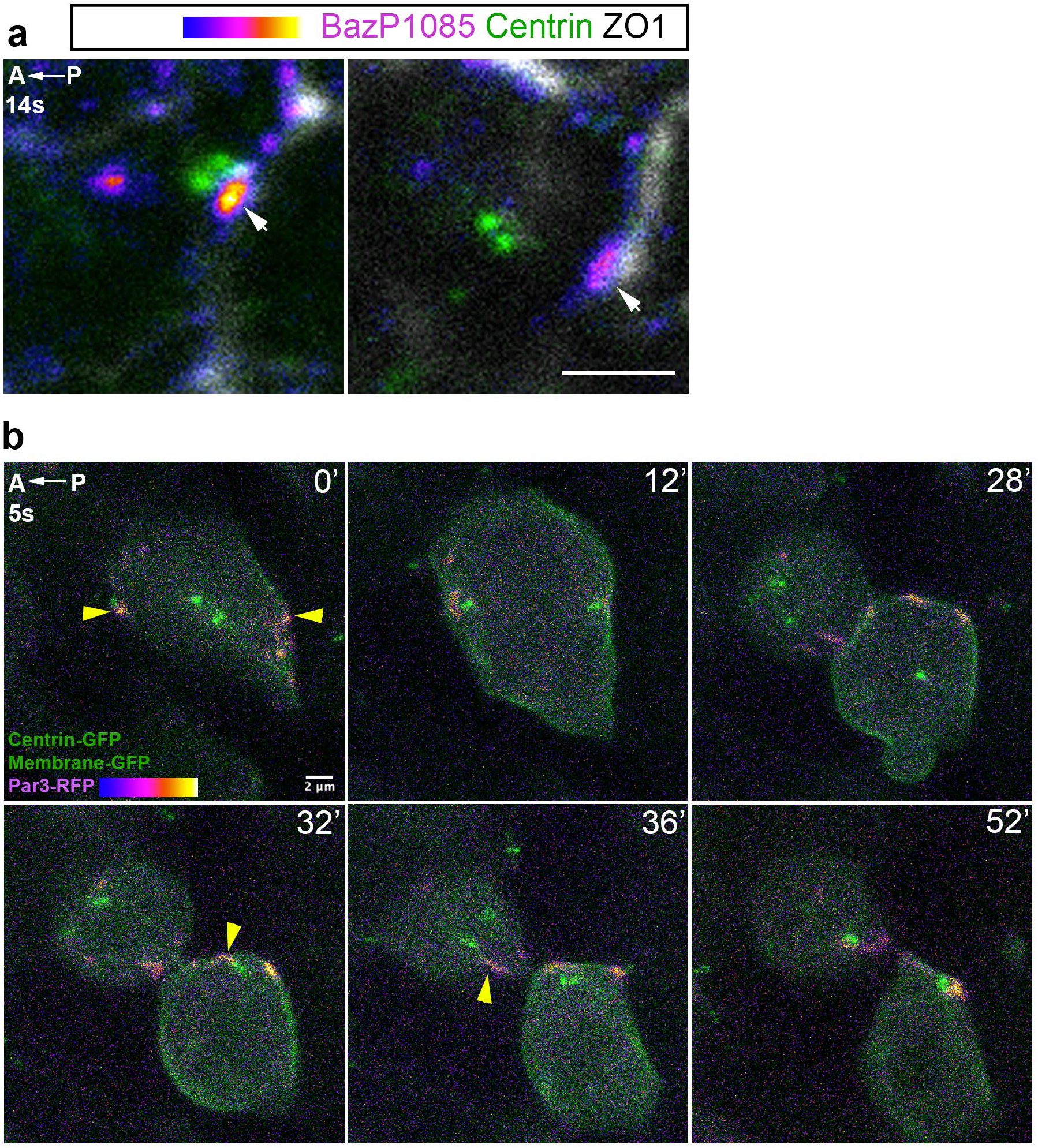
Par3 phosphorylation in FP cells and rapid migration of centrosome to Par3 patches after mitosis. **a)** Individual cells from dorsal views of 14 s stage embryos showing IF with an antibody against Drosophila Par3 (Bazooka/Baz) phosphorylated on Ser1085 (BazP1085) in FP cells. Two distinct cells are shown. phospho-Par3 displays the same patchy localization as total Par3 (Fig4a) White arrows point at patches at transverse membranes, which are present whether the BB is in contact with the posterior membrane (left images) or not (right image). **b)** Example of FP cell mitosis in an early stage (5s) embryo mosaically injected with Par3-RFP, Centrin-GFP and Membrane-GFP mRNAs and imaged every 4 minutes (MovieS12). Yellow arrows point at Par3-RFP patches, at the onset of mitosis (t=0’) and after cytokinesis, when the centrosome makes contact with the Par3 patch, in the posterior daughter cell (t=32’) and anterior daughter cell (t=36’).

**Supplementary Figure S4:**
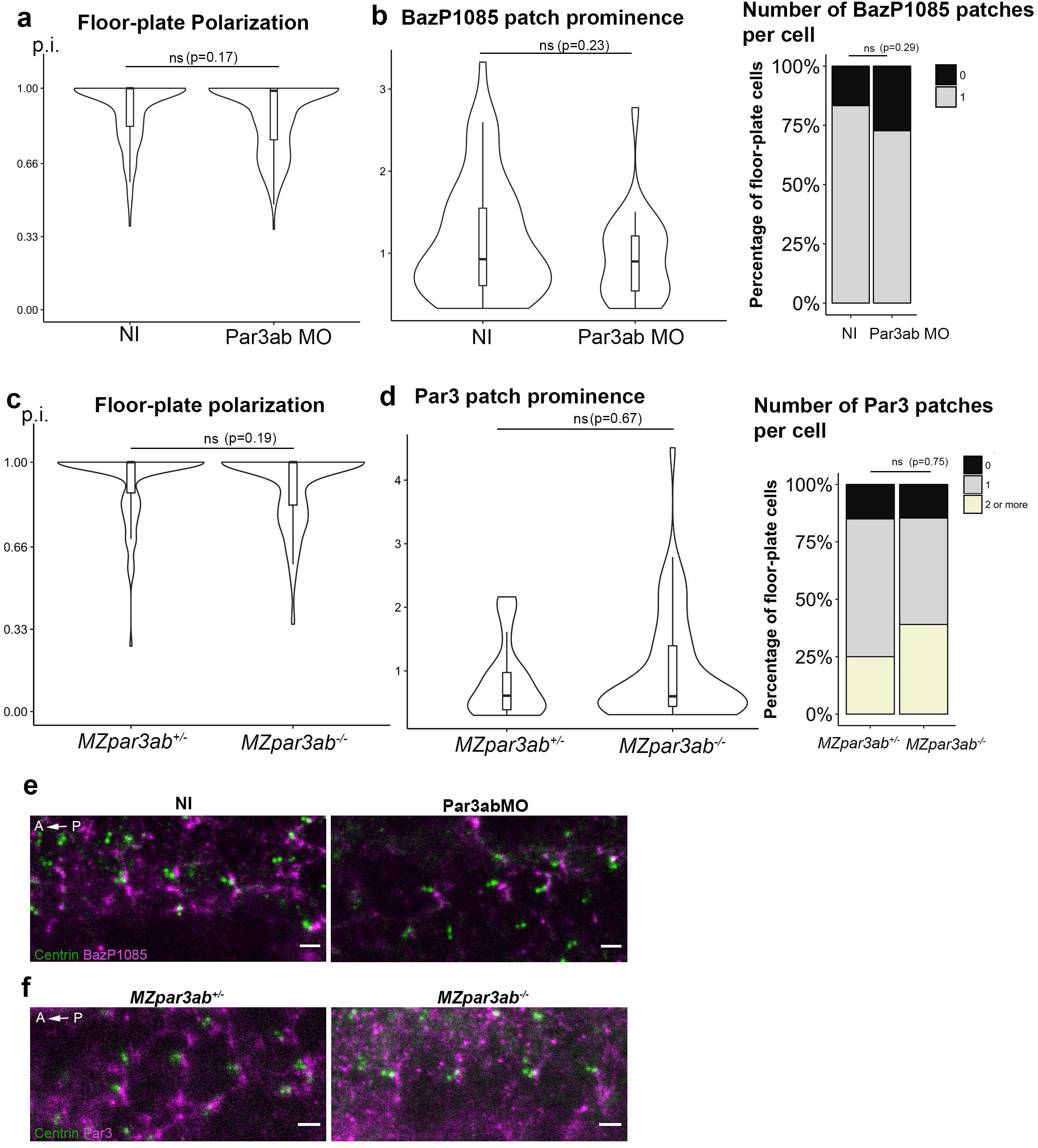
*Par3ab* morphants or mutants have normal FP polarization and Par3 patches. **a)** FP polarization index (p.i.) in non-injected (NI) and Par3ab morpholino (MO)- injected embryos at 18s stage (NI : 9 embryos, 171 cells; Par3ab MO : 16 embryos, 244 cells). **b)** BazP1085 patch prominence (left) and number (right) in NI and Par3ab MO injected embryos at 18s stage. NI : 4, embryos, 66 cells; Par3MO : 3 embryos, 38. cells **c)** p.i. of maternal zygotic heterozygous (*MZpar3ab^+/-^*) or homozygous (*Mpar3ab^-/-^*) *par3ab* mutants at 18s stage. *MZpar3ab^+/-^* : 7 embryos, 106 cells; *MZpar3ab^-/-^* : 9 embryos, 152 cells. **d)** Par3 patches prominence (left) and number (right) in maternal zygotic heterozygous (*MZpar3ab^+/-^*) or homozygous (*MZpar3ab^-/-^*) Par3ab mutants at 18s stage (*MZpar3ab^+/-^* : 3 embryos, 27 cells; *MZpar3ab^-/-^* : 3 embryos, 59 cells). p.i. and patch prominence are compared with a Wilcoxon test; Par3 or BazP1085 patches number are compared with Fisher test. **e)** Immunostaining of FP cells not injected (NI) or injected with Par3ab morpholino (Par3ab MO) showing the equivalent amount of BazP1085 staining in both conditions. **f)** Immunostaining of FP cells in *MZpar3ab^+/-^* and *MZpar3ab^-/-^* showing the equivalent amount of Par3 in both genotypes.

**Supplementary Figure S5:**
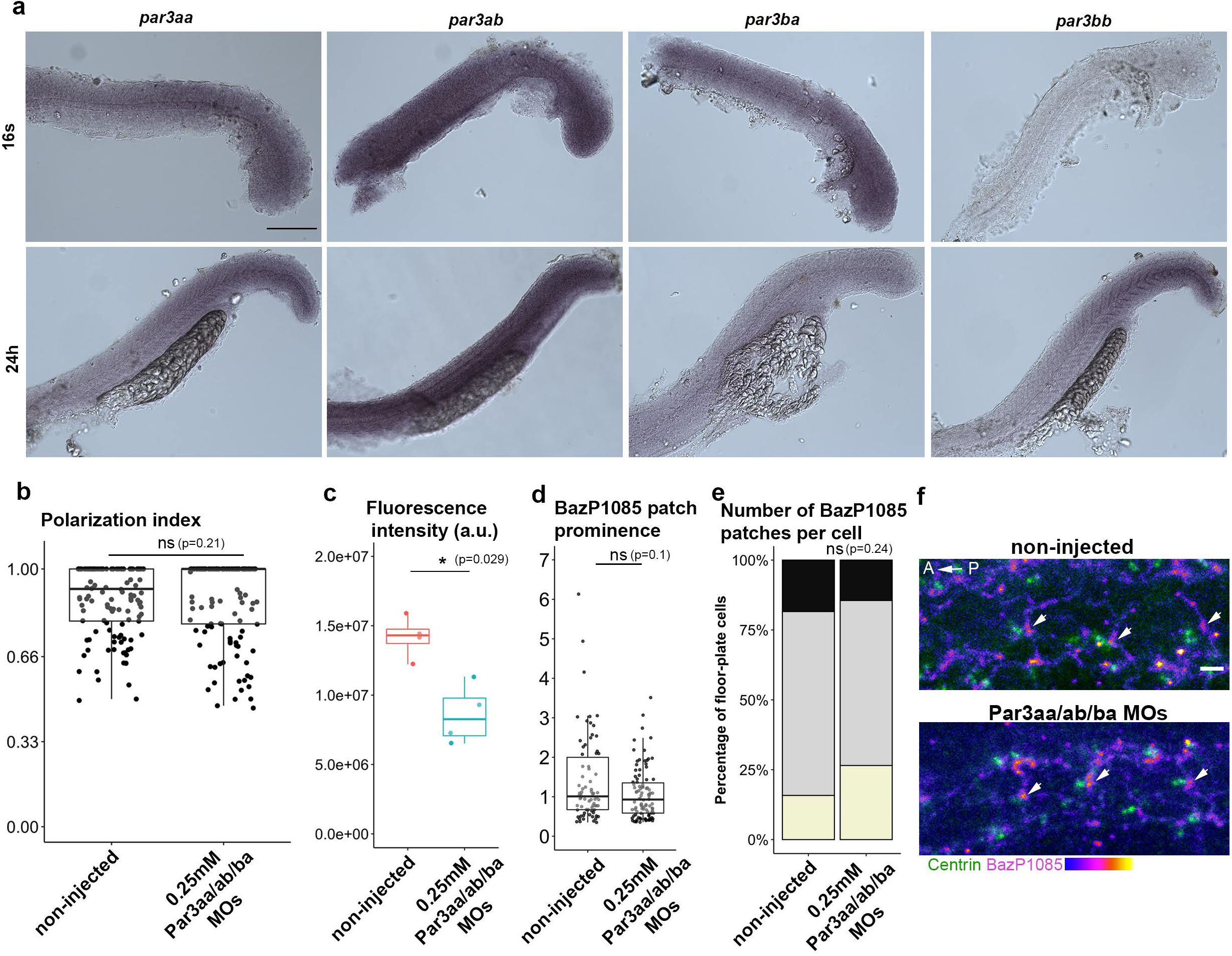
*par3aa/ab/ba* are broadly expressed during somitogenesis but their MO-mediated knockdown does not affect FP polarization. **a)** In situ hybridization reveals broad expression of *par3aa*, *ab* and *ba* genes at 16 s stage and 24 hpf. *par3bb* is not expressed at 16 s but is broadly expressed at 24 hpf with a posterior to anterior decreasing gradient. Scale bar 200µm. **b)** Polarization index of 18 s stage embryos FP after coinjection of 3 MOs targeting *par3aa*, *ab* and *ba* at the one cell stage (non-injected: 4 embryos, 264 cells; morphants: 4 embryos, 133 cells). **c)** Global phosphorylated-Par3 immunostaining signal (sum of Par3 fluorescence on a similar portion of the floor-plate, a.u. arbitrary unit, 4 embryos for each condition). **d)** Phospho-Par3 patches prominence in non-injected embryos and triple morphants **e)** Number of phospho-Par3 patches per cell in non-injected embryos and triple morphants **f)** Immunostaining of FP cells in non-injected and triple morphants showing the presence of phospho-Par3 patches (white arrows) in both conditions. c-e) non-injected: 3 embryos, 76 cells; morphants 4 embryos, 83 cells. Scale bar: 2µm b,c,d) Comparison done with a Wilcoxon test, e) Comparison done with a Fisher test

## SUPPLEMENTARY MOVIES LEGENDS

MovieS1: **Live imaging of a BB bouncing off the posterior membrane in an early-stage FP cell**.

wt embryos were injected with Centrin-GFP (green) and membrane-Cherry (magenta) mRNAs at the one-cell stage. White arrows indicate the position of the BB at the first and last time-points. Images were taken every 5 minutes during the 6 s to 9 s stages time-frame. Dorsal view. Corresponds to Fig2a.

MovieS2: **Live imaging of a BB bouncing off posterior and anterior membranes in an early-stage FP cell.**

wt embryos were injected with Centrin-GFP (green) and membrane-Cherry (magenta) mRNAs at the one-cell stage. White arrows indicate the position of the BB at the first and last time-points. Images were taken every 5 minutes during the 6 s to 9 s stages time-frame. Dorsal view. Corresponds to Fig2b.

MovieS3: **Live imaging of a BB staying in contact with the posterior membrane in a late-stage FP cell.**

wt embryos were injected with Centrin-GFP (green) and membrane-Cherry (magenta) mRNAs at the one-cell stage. White arrows indicate the position of the BB at the first and last time-points. Images were taken every 5 minutes during the 18 s to 21 s stages time-frame. Dorsal view. Corresponds to Fig2c.

MovieS4: **Live imaging of BB bouncing against the posterior membrane in a late-stage FP cell.**

wt embryos were injected with Centrin-GFP (green) and membrane-Cherry (magenta) mRNAs at the one-cell stage White arrows indicate the position of the BB at the first and last time-points. Images were taken every 5 minutes during the 18 s to 21 s stages time-frame. Dorsal view. Corresponds to Fig2d.

MovieS5: **Live imaging of BB movements in a FP cell displaying a membrane digitation between BB and the posterior membrane** (yellow arrow at t=115 min).

wt embryos were injected with Centrin-GFP (green) and membrane-Cherry (magenta) mRNAs at the one-cell stage. White arrows point at the BB. Images were taken every 5 minutes during the 6 s to 9 s stages time-frame. Dorsal view. Corresponds to Fig3a upper row.

MovieS6: **Live imaging of BB movements in a FP cell displaying a membrane digitation between BB and the anterior membrane** (yellow arrow at t=18 min).

wt embryos were injected with Centrin-GFP (green) and membrane-Cherry (magenta) mRNAs at the one-cell stage. Membrane digitations between the posterior membrane and BB can also be seen at t=10min, t=26min and t=66min. White arrows point at the BB. Images were taken every 2 minutes during the 8 s to 10 s stages time-frame. Dorsal view. Corresponds to Fig3a lower row.

MovieS7: **Live imaging of microtubule dynamics and BB movements in a FP cell at the 5s stage.**

wt embryos were injected with EB3-GFP, Centrin-GFP (Fire LUT, coding for fluorescence intensity) and membrane-Cherry (green) mRNAs at the 16-cell stage. and then imaged for 8min every 10 seconds at the 5s stage. White arrow at t=0’’ points at the BB. White arrows at t=150’’, t=160’’, t=280’’ and t=290’’ point at posterior membrane deformation and yellow arrowheads at t=150’’ and t=280’’ point at microtubules linking BB and posterior membrane at the spot where the membrane bends and toward which BB will move. Dorsal view. Corresponds to FigS2d.

MovieS8: **Live imaging of BB movements and Par3-RFP localization in a polarizing FP cell.**

wt embryos mosaically expressing Centrin-GFP (green) and Par3-RFP (magenta). White arrows point at the BB at t=0 and at t=30 min, when the BB touches the posterior membrane. Images were taken every 2 min during the 15 s to 17 s stages time-frame. Lateral view. Corresponds to Fig4c.

MovieS9: **Live imaging of BB movements and Par3-RFP localization in a non-polarizing FP cell.**

wt embryos mosaically expressing Centrin-GFP (green) and Par3-RFP (magenta). White arrows point at the BB at the beginning and end of movie. Images were taken every 5 minutes during the 17 s to 19 s stages time-frame. Lateral view. Corresponds to Fig4d.

MovieS10: **Live imaging of BB/Par3 patch contacts in an early-stage FP cell.**

wt embryo mosaically expressing Centrin-GFP, Membrane-GFP (green) and Par3-RFP (magenta). White arrows point at the BB at the beginning of the movie, when the BB is in contact with the anterior Par3 patch, at t=30 min when it makes a contact with the posterior Par3 patch and at the end of the movie. Images were taken every 2 min during the 4 s to 5 s stages time-frame. Dorsal view. Corresponds to the most anterior cell in Fig4e.

MovieS11: **Live imaging of membrane digitations at Par3 patches in early stage FP cells.**

wt embryo mosaically expressing Centrin-GFP, Membrane-GFP (green) and Par3-RFP (magenta). White arrows point at the BB at the beginning and at the end of the movie. Yellow arrows at t=0 and t=68 min point at membrane digitations originating from the posterior and the anterior Par3 patches, respectively. Images were taken every 4 min during the 7 s to 8 s stages time-frame. Dorsal view. Corresponds to Fig4f.

MovieS12: **Live imaging of BB moving to Par3 patches after cytokinesis in dividing early stage FP cells.**

wt embryo mosaically expressing Centrin-GFP, Membrane-GFP (green) and Par3-RFP (Fire LUT, coding for fluorescence intensity). Yellow arrows at t=0 point at the posterior and the anterior Par3 patches which are present in FP cells in interphase. Yellow arrows at t=32min and t=36min point at BB/Par3 patch contact in posterior and anterior daughter cells respectively. Images were taken every 4 min during the 5s to 6s stages. Dorsal view. Corresponds to FigS3b.

MovieS13: **Live imaging of BB/<u>lateral</u> Par3 patch contacts in an early-stage FP cell of a *vangl2^m209 / m209^* mutant.**

*vangl2^m209/m209^* embryo mosaically expressing Centrin-GFP, Membrane-GFP (green) and Par3-RFP (magenta). White arrows point at the BB at the beginning and at the end of the movie. Images were taken every 4 min during the 5 s to 6 s stages time-frame. Dorsal view. Corresponds to Fig7f.

